# Ubiquitin-like SUMO protease expansion in rice (*Oryza sativa*)

**DOI:** 10.1101/2025.08.20.671006

**Authors:** Kawinnat Sue-ob, Cunjin Zhang, Eshan Sharma, Rahul Bhosale, Ari Sadanandom, Andrew R Jones

**Affiliations:** Institute of Systems, Molecular and Integrative Biology, Department of Biochemistry, Cell and Systems Biology, University of Liverpool, Liverpool, L69 7ZB, United Kingdom; Department of Biosciences, Durham University, Durham, DH1 3LE, United Kingdom; School of Biosciences, University of Nottingham, Nottingham, LE12 5RD, United Kingdom

**Keywords:** *Oryza sativa*, rice pan-genome, SUMOylation, SUMO protease, Ubiquitin-like protease (ULP)

## Abstract

- SUMOylation is a protein post-translational modification that is essential for plant growth and response to changing environments. However, past work in this area has mainly focused on simple sequence similarity methods for discovering SUMOylation genes, often using orthologue mapping from yeast or *Arabidopsis*.
- In this work, we employed a range of computational techniques and approaches to describe and characterise the SUMOylation machinery in Asian rice (*Oryza sativa*), as a globally important stable crop, and where the SUMOylation system has been shown to play key roles in responses to biotic and abiotic stresses. We describe and analyse the ULP system at the phylogenetic, transcriptional, and protein structural levels, with a focus on the rice reference genome, a well-annotated Rice Population Reference Panel (RPRP), and wild rice genomes.
- Our analysis revealed the expansion of ULPs in the reference genome and RPRP set (32 – 45 ULPs) compared to wild rice (9-36 ULPs), raising an intriguing hypothesis about the expansion of the ULP family being driven by selective breeding pressure.
- We provide evidence of potential functional ULPs and their possible roles in biotic and abiotic stress responses in cultivated rice. These insights offer valuable resources for future rice breeding and crop improvement.

## Introduction

SUMOylation is a post-translational modification (PTM) found across eukaryotes. The mechanisms of SUMOylation are best understood in model organisms such as *Saccharomyces cerevisiae*, *Arabidopsis thaliana* (Roy & Sadanandom, 2021) and *Homo sapiens* (Celen & Sahin, 2020). The key protein of this mechanism is SUMO (Small Ubiquitin-like Modifier), a ∼100 amino acid (aa) protein with a conserved diglycine (-GG-) motif at the C-terminus. Pre-mature SUMO is first cleaved at the C-terminal tail to reveal the di-Glycine by a ULP (Ubiquitin-like protease). In *A. thaliana*, the mature SUMO is activated by a complex of SAE1a/b and SAE2 (SUMO activating enzymes), in an ATP-dependent process. The activated SUMO is then conjugated to SCE1 (E2 conjugating enzyme) and subsequently transferred and ligated to a lysine (K) residue on a target protein by SIZ1 or HPY1 (E3 ligases). The SUMO can also be polymerised by the function of PIAL1/2 (Protein Inhibitor of Activated STAT-Like 1/2). The effects of SUMOylation of a target include localisation, complex formation, interaction with other proteins, enzyme activities, or prevention from ubiquitination (Roy & Sadanandom, 2021). Finally, SUMO proteases such as the DeSI and ULP types can cleave the SUMO off the targets, allowing targets and the free SUMO to be recycled.

The SUMOylation system plays various regulatory roles in plant vegetative development and maturation stages at almost every gene regulation step, including chromatin modification/remodelling, epigenetics, transcriptional activation/repression, RNA processing, protein trafficking/localisation and enzyme activities (Augustine & Vierstra, 2018; Benlloch & Lois, 2018; Roy & Sadanandom, 2021). In rice, OsSIZ2, an E3 ligase has been shown to modulate developmental processes, phosphate (Pei et al., 2017) and nitrogen homeostasis (Pei et al., 2019; Wang et al., 2015) and play a role in reproduction (Pei et al., 2019). Among SUMO proteases in rice, OsFUG1 is involved in regulating development and fertility, OsELS1 contributes to the control of flowering time (Rosa et al., 2018), and OsOTS1 plays a role in seed germination and root development (Srivastava, Zhang, & Sadanandom, 2016).

Beyond development, SUMOylation also plays crucial roles in plant response to both abiotic and biotic stresses. Overexpression of OsSCE1 (E2) reduced tolerance to drought stress, and vice versa in OsSCE1-knocked-down plants (Nurdiani et al., 2018). OsSCE2/3 were up-regulated during drought treatment (Joo et al., 2019). OsSCE3, interestingly, also responded to Abscisic acid (ABA), Gibberellin (GA), glucose, and H_2_O_2_ treatments (Joo et al., 2019). The overexpression of OsSIZ1 (E3) in *Arabidopsis* enhances tolerance to several stresses, including drought, heat, and salt (Mishra et al., 2018). Overexpression of OsOTS1 (ULP) in salt stress treatments promotes root growth and results in enhanced salt tolerance (Srivastava et al., 2016), whereas depletion of OsOTS1/2 reduces salt tolerance but increases drought tolerance (Srivastava et al., 2017). For biotic defence mechanisms, proteomic analyses show that the level of SUMO-conjugated proteins was increased after pathogen attack, such as *Pseudomonas syringae* (Bailey et al., 2016; Ingole et al., 2021). Given its regulatory potential, the SUMOylation system has been proposed as a potential target for crop improvement (as reviewed in Rosa et al., 2019 and Clark et al., 2021).

Previous studies have attempted to identify SUMOylation genes in crops such as rice, *O. sativa* (Teramura et al., 2017; Rosa et al., 2018; Orosa et al., 2018; Garrido et al., 2018), maize*, Z. mays* (Augustine et al., 2016), tomato, *S. lycopersicum* (Augustine et al., 2016; Zhang et al., 2018), potato, *S. tuberosum* (Augustine et al., 2016; Ghimire et al., 2020), and soybean, *G. max* (Li et al., 2017), using sequence similarity-based bioinformatics tools. These studies typically query known SUMOylation protein sequences from *A. thaliana* and *S. cerevisiae* (yeast; review in Clark et al., 2022). Among monocots, SUMO components are best characterised in maize and rice (Clark et al., 2022).

In rice, by searching for homologous genes of *AtSUMO1* and *ScSMT3*, studies have identified six *OsSUMO* modifiers (Augustine et al., 2016; Rosa et al., 2018; Teramura et al., 2021). A more recent study has made a putative identification of *OsSUMO7* (Ibrahim et al., 2021). However, the protein sequence for putative *OsSUMO7* lacks a conserved di-Gly motif followed by a short C-terminal region, a hallmark of true SUMO modifiers. Currently, two E1 activating enzymes (OsSAE1 and OsSAE2), three E2 conjugating enzymes (OsSCE1, OsSCE2, and OsSCE3), and three E3 ligases (OsSIZ1, OsSIZ2, and OsHPY2) have been identified in rice but no E4 ligase gene have been reported (Rosa et al., 2018).

Unlike the other SUMO components, SUMO proteases have highly divergent sequences at the N-terminus, which limits the discovery using BLAST-based methods. Furthermore, as proteases, the catalytic domains (HxNCN for DeSI and HxDxC for ULPs) are crucial for deSUMOylation (Kurepa et al., 2003). By searching for ULPs with conserved catalytic domains in the RGAP rice database, previous studies have suggested there are 12-22 ULPs in rice (Garrido et al., 2018; Srivastava, Zhang, et al., 2016; Yates et al., 2016). However, only 7 OsULPs (OsFUG1, OsELS1, OsSPF1, OsELS2, OsOTS1, OsOTS2, and OsOTS3) have been experimentally validated for the protease function (Rosa et al., 2018; Srivastava, Zhang, & Sadanandom, 2016). An expansion of SUMO proteases, relative to *Arabidopsis*, was also observed in maize (Yates et al., 2016). There has been past speculation that expansion of ULPs in crops could have resulted from domestication and could give rise to target specificity during deSUMOylation (Srivastava et al., 2016; Garrido et al., 2018; Verma et al., 2018; Morrell and Sadanandom 2019). DeSIs, on the other hand, are a relatively recent discovery. They were first characterised in *Arabidopsis* (Orosa et al., 2018; Roy & Sadanandom, 2021; Seok et al., 2021) and more recently in *O. sativa* but without molecular functional validation (Lian et al., 2024).

Despite the importance and conservation of SUMOylation in plant systems, there are gaps in our knowledge regarding the SUMO components, and the molecular mechanisms are elusive in crops (Clark et al., 2022). Identifying SUMOylation-related genes is a crucial first step in studying SUMOylation in crops. In this study, we aimed to profile SUMOylation components in rice (*O. sativa*), as there is an apparent and unusually large expansion of ULP proteases in rice. We also wished to explore variation in the profile of ULPs across the rice pan-genome, now that there are leveraging newly available multiple high-quality genome assemblies and gene models available, in the so-called Rice Population Reference Panel (RPRP) genome set (Zhou et al., 2023). Furthermore, we performed computational validation using structural biology and assessed public transcriptomic data to provide strong evidence for the functionality of novel ULPs that were predicted.

## Methods

### Identification of ULPs in *O. sativa* Japonica

Our first analysis aimed at discovery of ULPs in the reference genome *O. sativa* Japonica Nipponbare (ISRGP). We queried well-characterised ULPs from *S. cerevisiae* and *A. thaliana* to the two rice genome annotations: RAP-DB (Sakai et al., 2013) and RGAP (previously called “MSU”, v7, Kawahara et al, 2023) using BLASTP and TBLASTN (blast+/2.9.0, (Camacho et al., 2009)). The BLAST results were filtered with e-values < 0.05. The significant results from BLAST were analysed for protein domains by InterProScan v5.66-98.0 (Jones et al., 2014) with its default parameters.

The sequences bearing the ULP domain (IPR003653), as identified by InterProScan (Jones et al., 2014) were subjected to alignment using MUSCLE v5 (https://hub.docker.com/r/pegi3s/muscle, López-Fernández et al., 2021). Specifically, sequences displaying alignment with known ULPs from yeast and *Arabidopsis*, particularly within the catalytic domain characterized by the HxDxC motif, were retained as ULP candidates. Proteins derived from the RAP-DB and RGAP gene models were aligned separately with known ULPs from yeast and *Arabidopsis*.

### ULP genes identification in the RPRP and wild rice population

We used BLASTP to search for the ULP orthologues within proteins annotated in the RPRP genomes: Aromatic (*Os*ARC), Indica (*Os*GoSa, *Os*IR64, *Os*KYG, *Os*LaMu, *Os*Lima, *Os*LiXu, *Os*MH63, *Os*Pr106, and *Os*ZS97), Aus (*Os*N22, *Os*NaBo), Tropical Japonica or Tropjap (*Os*Azu, *Os*CMeo, *Os*KeNa), Temperate Japonica or Tempjap (IRGSP RefSeq – three annotation sets RAP-DB, RGAP and a new annotation set from the Gramene database, called *Os*Nip), and in wild rice: *O. brachyantha*, *O. glumipatula*, *O. longistaminata*, *O. nivara*, *O. punctata, O. rufipogon*, *O. barthii*, *O. glaberrima* and closely rice related grass; *L. perrieri* (see the genome information in **Table S1**) by querying 37 ULP protein sequences from the previous step (both RAP-DB and RGAP models, **Table S2**). A stringent filter criterion (e-value < 10^−6^) was applied to BLASTP results, given that searches were performed within genus level (Oryza), we would expect orthologues to have low e-values. Significant (protein) matches resulting from BLASTP were next analysed with InterProScan. Those with a significant match to the ULP domain (IPR003653) were then aligned with the ULP reference proteins from yeast and *Arabidopsis*.

For the RPRP dataset, with the information of pan-gene set from the “PanOryza” pan-gene project (Contreras-Moreira et al., 2025), we were able to map all the ULP candidates to the pan-gene matrix and investigate at the “pan-gene” levels. Here a pan-gene is cluster of genes that have overlapping coordinates following whole genome alignment, which ideally correspond to alleles of the same gene. If gene models have been incorrectly called in some aligned genomes, then a pan-gene can contain predicted transcript / protein sequences that differ significantly. Protein sequences derived from each pan-gene were then independently aligned with yeast and *Arabidopsis* ULP proteins (i.e. one alignment per pan-gene) using three different multiple sequence alignment (MSA) algorithms - MUSCLE, MAFFT, and CLUSTALOmega (https://hub.docker.com/r/pegi3s/muscle, https://hub.docker.com/r/ pegi3s/mafft, and https://hub.docker.com/r/pegi3s/clustalomega, docker images (López-Fernández et al., 2022)) to optimise the identification of conserved ULP active sites. If results from MSAs differed in the aligned residues at the known active sites (from yeast and *Arabidopsis*), we selected if a given protein had the required motifs aligned in at least one set of alignment results, then it was called as a correct match to go forward for the next analysis step. For wild rice species, we aligned all ULP candidate proteins (from BLASTP and InterProScan) with known yeast and *Arabidopsis* ULPs from each wild rice species at a time.

### Phylogenetic analysis

A phylogenetic tree was generated, derived from protein MSAs for *S.cerevisiae*, *A.thaliana*, *L. perrieri*, wild rice, and RPRP, by the Neighbour-joining method with 1000 bootstraps in MEGA X (Kumar et al., 2018) and visualised/annotated in iTOL (Letunic & Bork, 2024). The 50 sub-clusters were derived from cutree function in ape library (Paradis & Schliep, 2019). The Yeast ULP1 (ScULP1) was used as the outgroup.

### Selection of representative genes from the RPRP pan-genes

To select a representative ULP gene from within the pan-genes from the RPRP set, we selected a confident ULP candidate (i.e. containing the ULP domain and correct catalytic sites) following the order of genomes/annotations sets within the pan-gene matrix (Contreras-Moreira et al., 2025) i.e. RAP-DB, OsNip (Gramene gene model), and RGAP (from the IRGSP Nipponbare reference genome), respectively. If the pan-gene did not contain a Nipponbare gene member, or those members did not pass the ULP identification criteria, a representative was picked in the following order: OsAzu, OsCMeo, OsPr106, OsKeNa, OsARC, OsZS97, OsN22, OsMH63, OsNaBo, OsLiXu, OsGoSa, OsLaMu, OsKYG, OsIR64, and OsLima (Contreras-Moreira et al., 2025).

### Protein disorder region prediction

We utilized metapredict v2.61 (Emenecker et al., 2021) to identify disordered regions within protein sequences. The representative ULP sequences from pan-gene clusters and all ULPs from the eight wide rice varieties, were input into metapredict, and the results were visualized using R.

### Protein structure analysis of the ULPs

Proteins structures for the ULP representatives from the RPRP pan-genes and all identified ULPs in wild rice were predicted using colabfold (https://github.com/YoshitakaMo/localcolabfold; Mirdita et al. (2022)). The colabfold parameters were set as generating three models (--num-models 3), using amber for the structure refinement (--amber) and running on GPU (--use-gpu-relax). Only the relaxed_rank_001 structures were further investigated. We measured the distances between the three amino acids at the catalytic site (angstrom, Å) using in-house python script (and BioPython version 1.81), and visualised using PyMOL 2.1.1 (Schrodinger, 2015).

### RNA-sequencing analysis

Public rice RNA-sequencing datasets were collected and re-analysed by Yu et al., (2022) using a uniform pipeline. Briefly, the fastq files were mapped to RGAP genome annotation using HISAT2 (Kim et al., 2019). The duplicated reads were removed by using SAMtools (rmdup) and the transcripts abundance (Fragments Per Kilobase of transcript per Million mapped reads; FPKM) were calculated using StringTie (Pertea et al., 2015). FPKM values of ULPs (RGAP models) were derived from Public RNA-seq Database (https://plantrnadb.com/ricerna/), scaled (z-score), clustered by default parameters (complete method and Euclidean distance) and visualised using ComplexHeatmap library in R (Gu et al., 2016).

## Results

### Identification of ULPs in the O. sativa Nipponbare reference genome

To identify ULPs in the *O. sativa* IRGSP (Nipponbare) Reference Sequence (hereafter IRGSP RefSeq), we performed a two-step identification approach. Firstly, homologs of the two yeast ULPs (ScULP1 and ScULP2) and eight *Arabidopsis’s* ULPs (AtOTS1, AtOTS2, AtELS1, AtELS2, AtEDS4, AtSPF1, AtSPF2, and AtFUG1) were searched (using BLASTP and TBLASTN) against the two IRGSP RefSeq annotation sets (RAP-DB and RGAP). Significant results (e-value < 0.05) were identified and then analysed for protein domains using InterProScan. After performing the multiple sequence alignment (MSA), we carried forward and retained only those proteins from RAP-DB or RGAP gene models that had the catalytic domain (HxDxC) when aligned with the ten known ULPs from yeast and *Arabidopsis*. The alignment at the conserved positions catalytic motif (HxDxC) distinguishes ULPs from other proteases in the DUB (deubiquitination) family, such as UBPs (ubiquitination) (**Fig. S1**). Our analysis identified 33 ULPs (RAP-DB) or 37 (RGAP) and non-redundant models 37 ULPs (non-redundant models) in IRGSP RefSeq that contain the catalytic domains (**Table S2**). This indicates the inconsistency between the two gene models and highlights the importance of cross-referencing the genes of interest in both models, which should always be considered.

Secondly, the 37 candidates from the genes identified in the previous step (**Table S2**) were searched against a recently released additional gene model annotation set for IRGSP, recently produced and released by the Gramene database team (gene id starts with OsNip) using a similar approach, mapped to the pan-gene matrix and investigated at the pan-gene level (see Methods). Next, we further validated the candidates by predicting their 3D structures of the ULP proteins using Colabfold (Mirdita et al., 2022). Finally, the proteins that have conserved catalytic sites with yeast and *Arabidopsis* were finalised as ULPs in IRGSP RefSeq (**Table 1**, **Fig. 1**).

**Fig. 1.**
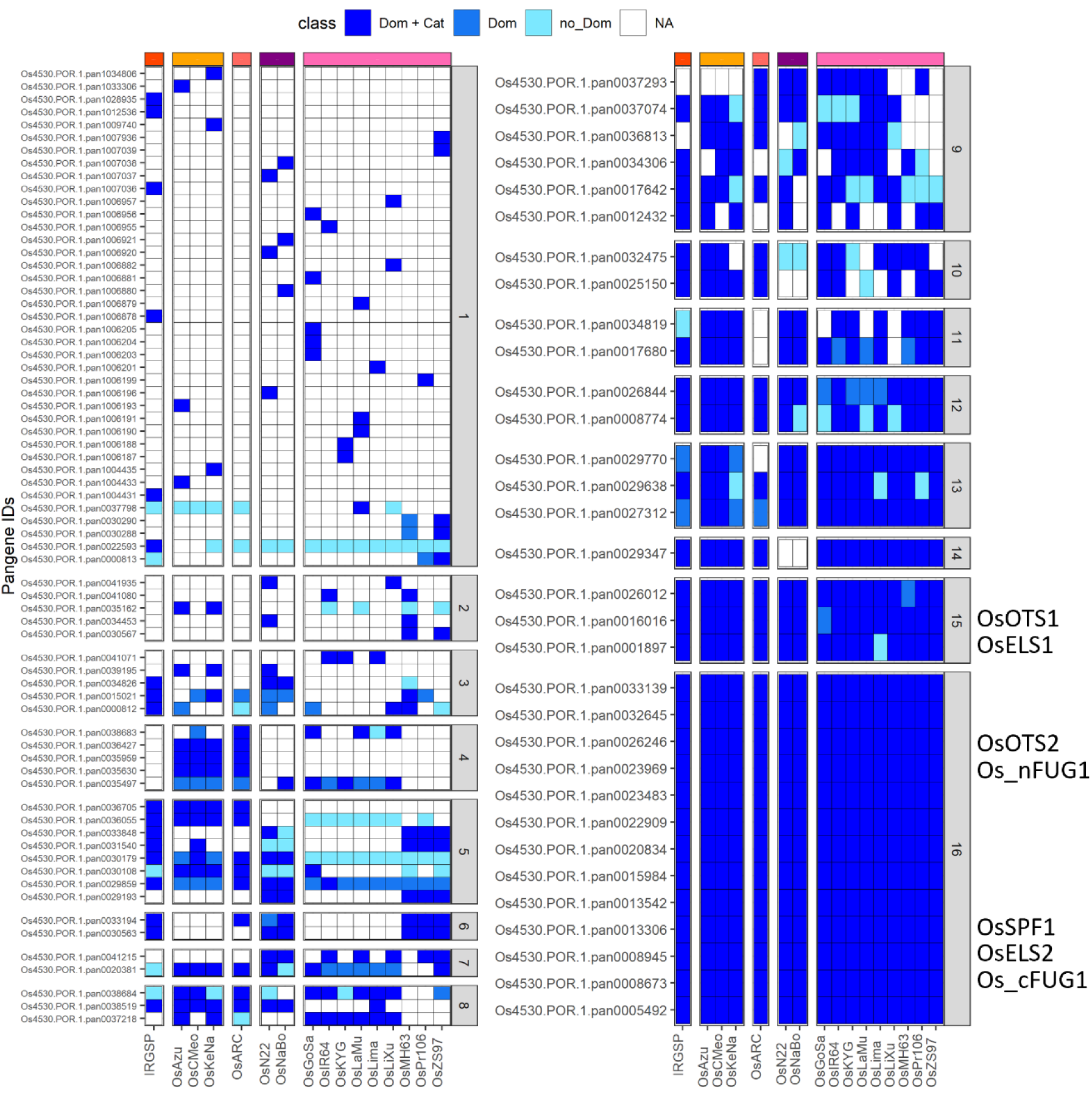
Occupancy of ULP-domain in RPRP genomes. 101 Pan-gene ids of ULP genes are shown in rows, and the groups of RPRP rice are separated by columns (Tempjap, Tropjap, Aro, Aus, and Indica, respectively). The pan-gene clusters are separated by the presence of ULP domains amongst 16 populations (domain occupancy). Colours show classification of the genes, ranging from the darkest blue to white if there are ULP domains and correct catalytic sites (Dom + Cat), only ULP domains (Dom), no ULP domains (no_Dom), and no genes (NA) in the pan-gene cluster.

**Table 1.** Summary of ULPs in *O. sativa* Japonica. The pan-gene identifiers (Os4530.POR.1.panXXXXXXX) of ULPs in IRGSP RefSeq (Nipponbare_merged) are arranged by the number of domain occupancy (count-domain) and genome occupancy (count-rice) in the RPRP set. Gene models containing a Ulp domain, as identified by InterProScan, are italicised; those with HxDxC catalytic residues, identified by MSA, are underlined; and the representative models that were protein-structured (Alphafold) are in bold. The gene names were assigned based on the hierarchical clusters in **Fig. 2** and followed by letters according to domain occupancy.

| Pan-id | RAP-db | Gramene | RGAP (MSU) | Gene name | count domain | count rice |
| --- | --- | --- | --- | --- | --- | --- |
| pan0005492 | **<u>Os02g0472200</u>** | None | <u>LOC_Os02g27280</u> | *OsULP40a* | 16 | 16 |
| pan0008673 | **<u>Os03g0344300</u>** | <u>OsNip_03G017390</u> | <u>LOC_Os03g22400</u> | (*OsFUG1*)<br>*OsELS3* | 16 | 16 |
| pan0008945 | Os03g0410100 | **<u>OsNip_03G021330</u>** | <u>LOC_Os03g29630</u> | *OsELS2* | 16 | 16 |
| pan0013306 | **<u>Os05g0207900</u>** | <u>OsNip_05G007210</u> | <u>LOC_Os05g11770</u> | *OsSPF1* | 16 | 16 |
| pan0013542 | **<u>Os05g0309000</u>** | <u>OsNip_05G011330</u> | None | *OsULP40b* | 16 | 16 |
| pan0015984 | **<u>Os06g0474600</u>** | <u>OsNip_06G015830</u> | <u>LOC_Os06g28030</u> | *OsULP40c* | 16 | 16 |
| pan0020834 | Os09g0258700 | **<u>OsNip_09G002660</u>** | <u>LOC_Os09g08440</u> | *OsULP50a* | 16 | 16 |
| pan0032645 | None | **<u>OsNip_09G004490</u>** | <u>LOC_Os09g12480</u> | *OsULP45a* | 16 | 16 |
| pan0033139 | Os10g0157000 | **<u>OsNip_10G003120</u>** | <u>LOC_Os10g06910</u> | *OsULP42a* | 16 | 16 |
| pan0022909 | **<u>Os10g0474000</u>** | <u>OsNip_10G014370</u> | <u>LOC_Os10g33450</u> | *OsULP35* | 16 | 16 |
| pan0023483 | **<u>Os11g0102700</u>** | <u>OsNip_11G000130</u> | <u>LOC_Os11g01180</u> | *OsULP37a* | 16 | 16 |
| pan0023969 | **<u>Os11g0214001</u>** | <u>OsNip_11G007290</u> | <u>LOC_Os11g10780</u> | *OsFUG1* | 16 | 16 |
| pan0026246 | **<u>Os12g0606900</u>** | <u>OsNip_12G021260</u> | <u>LOC_Os12g41380</u> | *OsOTS2* | 16 | 16 |
| pan0001897 | **<u>Os01g0355900</u>** | <u>OsNip_01G016190</u> | <u>LOC_Os01g25370</u> | *OsELS1* | 15 | 16 |
| pan0016016 | **<u>Os06g0487900</u>** | <u>OsNip_06G016450</u> | <u>LOC_Os06g29310</u> | *OsOTS1* | 15 | 16 |
| pan0026012 | **<u>Os12g0551900</u>** | <u>OsNip_12G017880</u> | <u>LOC_Os12g36580</u> | *OsULP40d* | 15 | 16 |
| pan0029347 | Os04g0354300 | **<u>OsNip_04G010250</u>** | <u>LOC_Os04g28590</u> | *OsULP43a* | 14 | 14 |
| pan0029638 | Os04g0583400 | **<u>OsNip_04G025050</u>** | <u>LOC_Os04g49394</u> | *OsULP50b* | 13 | 16 |
| pan0026844 | None | **<u>OsNip_01G019120</u>** | <u>LOC_Os01g33520</u> | *OsULP32* | 12 | 16 |
| pan0008774 | Os03g0365250 | **<u>OsNip_03G018740</u>** | <u>LOC_Os03g24960</u><br><u>LOC_Os03g24980</u><br><u>LOC_Os03g25000</u> | *OsULP50c* | 12 | 16 |
| pan0017680 | Os07g0283600 | **<u>OsNip_07G010740</u>** | <u>LOC_Os07g18300</u> | *OsULP38* | 11 | 14 |
| pan0032475 | None | **<u>OsNip_09G000230</u>** | None | *OsULP46* | 10 | 13 |
| pan0025150 | **<u>Os12g0127600</u>** | <u>OsNip_12G001960</u> | None | *OsULP37b* | 10 | 11 |
| pan0012432 | **<u>Os04g0639700</u>** | **<u>OsNip_04G028960</u>** | **<u>LOC_Os04g54670</u>** | **<u>OsULP44a</u>** | 9 | 9 |
| pan0037074 | None | **<u>OsNip_07G008460</u>** | **<u>LOC_Os07g12990</u>** | **<u>OsULP50d</u>** | 9 | 13 |
| pan0017642 | **<u>Os07g0267400</u>** | **<u>OsNip_07G010020</u>** | None | **<u>OsULP40e</u>** | 9 | 15 |
| pan0034306 | **<u>Os11g0646232</u>** | **<u>OsNip_11G022650</u>** | **<u>LOC_Os11g42610</u>** | **<u>OsULP40f</u>** | 9 | 11 |
| pan0038519 | **<u>Os12g0103580</u>** | **<u>OsNip_12G000230</u>** | **<u>LOC_Os12g01290</u>** | **<u>OsULP36</u>** | 8 | 8 |
| pan0033194 | **<u>Os04g0377400</u>** | **<u>OsNip_04G011280</u>** | **<u>LOC_Os04g30860</u>** | **<u>OsULP50e</u>** | 6 | 7 |
| pan0030563 | None | **<u>OsNip_06G009810</u>** | **<u>LOC_Os06g13750</u>** | **<u>OsULP48</u>** | 6 | 6 |
| pan0036055 | **<u>Os03g0614100</u>** | **<u>OsNip_03G026210</u>** | **<u>LOC_Os03g41780</u>** | **<u>OsULP42b</u>** | 5 | 12 |
| pan0029859 | None | **<u>OsNip_05G006220</u>** | **<u>LOC_Os05g10270</u>** | **<u>OsULP50f</u>** | 5 | 16 |
| pan0030179 | **<u>Os05g0417800</u>** | **<u>OsNip_05G017400</u>** | **<u>LOC_Os05g34520</u>** | **<u>OsULP45b</u>** | 5 | 16 |
| pan0036705 | **<u>Os05g0448000</u>** | **<u>OsNip_05G019120</u>** | **<u>LOC_Os05g37550</u>** | **<u>OsULP43b</u>** | 5 | 5 |
| pan0031540 | **<u>Os07g0504300</u>** | **<u>OsNip_07G016450</u>** | **<u>LOC_Os07g32090</u>** | **<u>OsULP43c</u>** | 5 | 7 |
| pan0033848 | None | **<u>OsNip_11G008380</u>** | **<u>LOC_Os11g12500</u>** | **<u>OsULP50g</u>** | 5 | 6 |
| pan0000812 | **<u>Os01g0113000</u>** | **<u>OsNip_01G001010</u>** | **<u>LOC_Os01g02270</u>** | **<u>OsULP44b</u>** | 3 | 8 |
| pan0015021 | **<u>Os06g0122600</u>** | **<u>OsNip_06G001600</u>** | **<u>LOC_Os06g03180</u>** | **<u>OsULP44c</u>** | 3 | 8 |
| pan0034826 | None | **<u>OsNip_12G012470</u>** | **<u>LOC_Os12g24880</u>** | **<u>OsULP19</u>** | 3 | 4 |
| pan1012536 | **<u>Os02g0262500</u>** | None | **<u>LOC_Os02g16240</u>** | **<u>OsULP28</u>** | 1 | 1 |
| pan1007036 | None | **<u>OsNip_03G004800</u>** | None | **<u>OsULP30</u>** | 1 | 1 |
| pan1006878 | **<u>Os03g0365200</u>** | None | **<u>LOC_Os03g24990</u>** | **<u>OsULP50h</u>** | 1 | 1 |
| pan1004431 | **<u>Os08g0429600</u>** | **<u>OsNip_08G017070</u>** | **<u>LOC_Os08g33280</u>** | **<u>OsULP50i</u>** | 1 | 1 |
| pan0022593 | **<u>Os10g0388600</u>** | **<u>OsNip_10G009270</u>** | **<u>LOC_Os10g24954</u>** | **<u>OsULP43d</u>** | 1 | 14 |
| pan1028935 | **<u>Os11g0235825</u>** | **<u>OsNip_11G022640</u>** | **<u>LOC_Os11g12780</u>** | **<u>OsULP39</u>** | 1 | 1 |

In total, we identified 45 ULPs in IRGSP RefSeq (**Table 1**, **Fig. 1**). In this final ULPs list, 12 genes, from the initial set (37 genes, RAP-DB/RGAP) were removed because 6 protein sequences were unaligned with known catalytic sites (when aligned within the pan-gene cluster) and 6 sequences have showed incorrect protein structures at catalytic sites, as predicted by Colabfold (**see Table S2**). In addition, an extra 20 genes were incorporated, indicating these proteins as their sequences were aligned better within their pan-gene cluster. Our approaches have successfully identified all previously known ULPs in rice, including OsOTS1 (Os06g0487900), OsOTS2 (Os12g0606900), OsFUG1 (Os03g22440), OsELS1 (Os01g25370), OsSPF1 (Os05g11770), and OsELS2 (Os03g29630), as previously described (Rosa et al., 2018). However, OsOTS3 (Os01g53630), which has no aligned catalytic sites after MSA, was discarded from the full gene final set. There are 13 ULPs that were found to contain the conserved ULP domains and correctly positioned catalytic sites across all 16 genomes examined (count_domain) (**Table 1**, **Fig. 1**).

### ULPs are expanded in the RPRP pan-genome

To further explore the expansion of ULPs in the rice pan-proteome (derived from the *RPRP* pan-genome), we identified ULPs from each RPRP genome/proteome using BLASTP, followed by protein domain identification (InterProScan) and MSA. As the identification of ULP relies on the MSA of the catalytic sites, there is, however, uncertainty of MSA that different algorithms produce different results (Blackburne & Whelan, 2013). To minimize these effects, we used MSA by using three different alignment algorithms to align for the ULPs (see method). Then, the identified ULP candidates were mapped to the pan-gene matrix. The pan-gene matrix was created via the GET_PANGENES pipelines (Contreras-Moreira et al., 2023), using the RPRP genomes and gene models as input, to identify the orthologous gene (pan-gene) in each genome. In total, the ULPs can be mapped to 101 pan-gene identifiers (after protein structure correction), with “genome occupancy” (**Fig. S3**) and “domain occupancy” (**Fig. 1**) ranging from 1-16 across the rice species. The ULPs have an unusual pattern of evolution – with strong enrichment in genes found in only a single genome. The analysis in Contreras-Moreira et al., 2025 described an approach for classifying protein function according to presence of a given domain across 16 genomes, with those protein domains with mean occupancy <10 (out of 16) being classed as “Highly variable” domains – which are enriched for functions related to disease resistance (amongst others), and expansion of the gene family appears to be driven by selection pressure. Based on the analysis here, the mean occupancy for the ULP is 5.94, raising the possibility that ULPs have also been under selection pressure.

Among the pan-gene clusters that have 16 genome occupancy in across all 16 rice species (27 clusters, **Fig. S3**), some of them were not identified as ULPs due to the absence of ULP domains or catalytic sites. The 16 domain occupancy group consists of 13 pan-gene clusters, including known rice ULPs: OsOTS2, OsSPF1, OsELS2, and Os_cFUG1 (**Fig. 1**). Two additional known rice ULPs, OsOTS1 and OsELS1, are grouped within the 15-domain occupancy cluster. This is because OsLima_OTS1 contains a ULP domain detected by InterProScan, but its catalytic sites do not align with the ULPs from yeast and *Arabidopsis,* while OsGoSa_ELS1 lacks the ULP domain (**Fig. S4**).

### ULP expansion may be associated with selected breeding in Asian rice

To examine whether the ULP expansion occurs only in cultivated rice species, we applied the same approach (BLASTP, followed by protein domain identification by InterProScan, 3-MSA, and protein structure modelling) to identify ULP encoding genes in eight wild rice species and compared the results with the RPRP pan-genome.

In wild rice, the number of ULPs ranges from 9 to 26, except for *O. punctata*, which contains 36 genes (**Fig. 2a**). We found that the counts of ULPs in the RPRP genomes were higher than in almost all wild rice species, with the exception of *O. punctata* (**Fig. 2a**). Next, we constructed the phylogenetic tree (using Neighbour joining method with 1000 bootstraps) to show the relationship among known ULPs in yeast, *Arabidopsis*, and newly identified ULPs in cutgrass, wild and domesticated rice (RPRP set) (**Fig. 2b**). The dendrogram was then divided into 50 clusters of ULPs (**Table S3**). There are clusters containing known rice ULPs: OsOTS1 (Os06g0487900 or LOC_Os06g29310; cluster 11), OsOTS2 (Os12g0606900 or LOC_Os12g41380; cluster 12), OsSPF1 (Os05g0207900 or LOC_Os05g11770; cluster 8). Additionally, OsELS1 (Os01g0355900 or LOC_OS01G25370), OsELS2 (Os03g0410100 or LOC_Os03g29630), and the currently designated OsFUG1 (Os03g0344300 or LOC_Os03g22400; Os_cFUG1) were assigned to cluster 24 – 26, respectively (**Fig. 2b**).

**Fig. 2.**
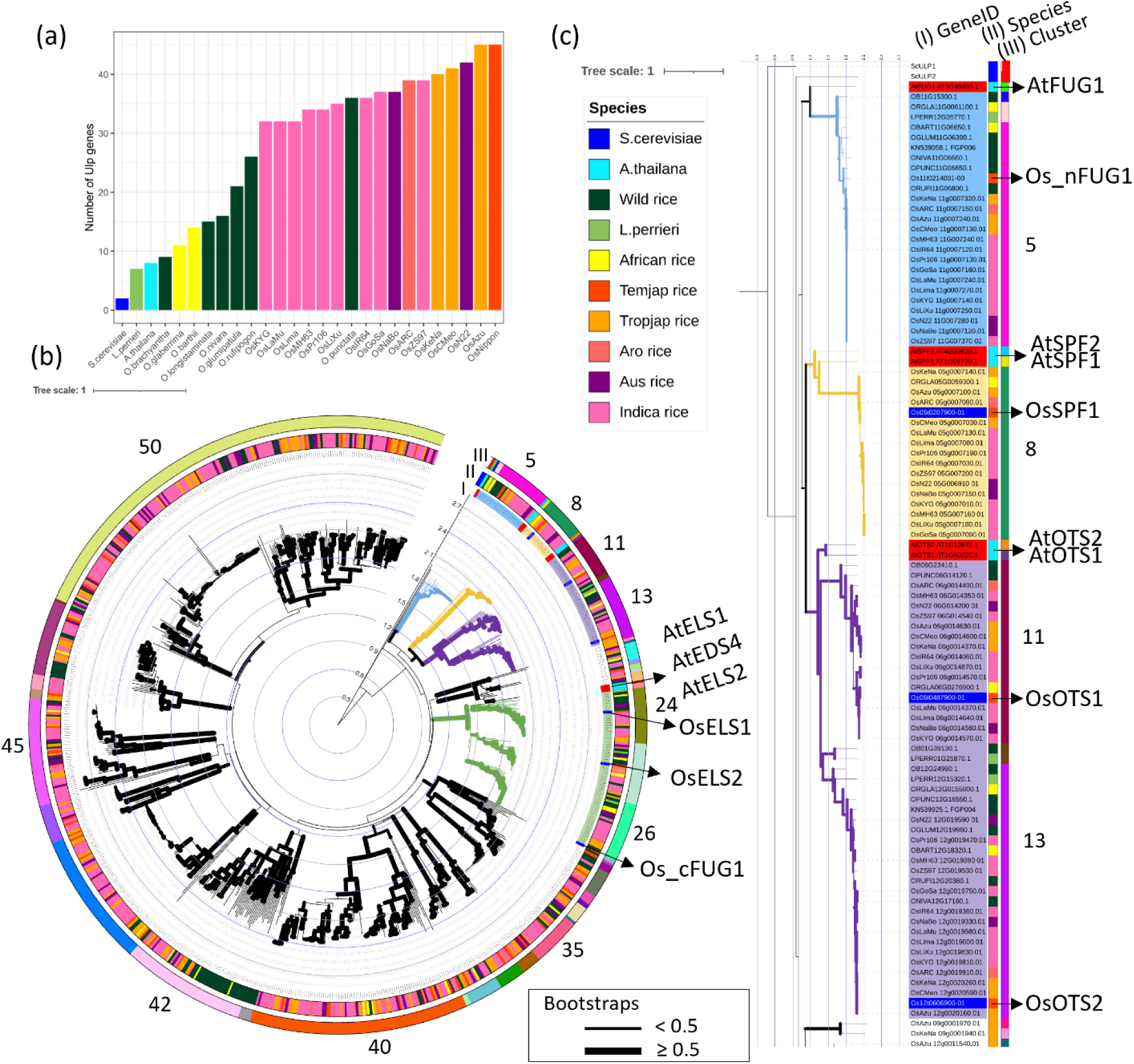
Expansion of ULP in rice. **a**) Count of ULP encoding genes (ranked by count) in yeast, *Arabidopsis*, cutgrass (*L. perrieri*), wild rice (*O. brachyantha*, *O. glumipatula*, *O. longistaminata*, *O. nivara*, *O. punctata, O. rufipogon*), African rice (*O. barthii*, *O. glaberrima*) and *O. sativa* RPRP genome set: Aro (*Os*ARC), Indica (*Os*GoSa, *Os*IR64, *Os*KYG, *Os*LaMu, *Os*Lima, *Os*LiXu, *Os*MH63, *Os*Pr106, and *Os*ZS97), Aus (*Os*N22, *Os*NaBo), Tropical Japonica or Tropjap (*Os*Azu, *Os*CMeo, *Os*KeNa), Temperate Japonica or Tempjap (ISRGP RefSeq). **b**) ULP tree of all species. The tree was constructed by Neighbour Joining (NJ) with 1000 bootstraps. Annotation strips show (I) gene ids, (II) species, (III) subclusters as also represent by numbers. See the order of ULPs of the phylogenetic tree in **Table S3**. **c)** Sub-cluster of ULPs (clusters 1-15), showing a potential novel OsFUG1 (Os_nFUG1) and the known ULPs in rice: OsOTS1, OsOTS2. The known ULPs in *Arabidopsis* and rice are highlighted in red and blue, respectively. The bootstrapping values over or equal to 50% are shown in thicker clades. The tree was rooted by using yeast ULP1a as an outgroup.

Interestingly, we identified a novel rice ULP group within cluster 5 (**Fig. 2b-2c**), which is grouped closely related to *A. thaliana* FUG1 (FOURTH ULP GENE CLASS 1). The role of AtFUG1 has been recently revealed as involving in epigenetic gene silencing (Sureshkumar et al., 2024). In rice, OsFUG1 was discovered by a homology-based search against RGAP reference genome (Srivastava et al., 2016) and was named OsFUG1 based on a phylogenetic relationship with AtFUG1 (Rosa et al., 2018). However, our analysis suggests that the current OsFUG1 (here called Os_cFUG1) is not a close homologue of AtFUG1, as Os_cFUG1 sits within cluster 26. Within cluster 5, which contains AtFUG1, we see that identified Os11g0214001 is a close homologue, thus making it a more appropriate candidate for renaming as OsFUG1 – here designated Os_nFUG1. Cluster 26, which includes Os_cFUG1, also contains ELS-type ULPs. ELS ULPs were initially characterised based on molecular and phenotypic approaches and have been implicated in flowering (Rosa et al., 2018). The close homology between Os_cFUG1 and OsELS1 / OsELS2 suggests that Os_cFUG1 should be renamed to OsELS3. Moreover, MSA shows that the AtFUG1 is more similar to Os_nFUG1 than Os_cFUG1 (Figure 2.S5. Additionally, there are ULPs in clusters 27 −50 that contain ULPs with no closely homologous genes in yeast and or *Arabidopsis*, suggesting that they could be novel groups of ULPs in rice (**Fig. 2b**).

Asian rice species were domesticated from two wild rice species, *O. rufipogon* and *O. nivara* (Jing et al., 2023). Among rice ULPs, *Os_cFUG1* (*Os03g0344300* or *LOC_Os03g22400*) has been identified as one of the domesticated genes that originated from *O. nivara* (Jing et al., 2023). However, our phylogenetic analysis suggests that OsFUG1 (Os_cFUG1) forms a sister clade with ORUFI03G17620 and is more distantly related to ONIVA03G18050, indicating that Os_cFUG1 is likely derived from *O. rufipogon* (cluster 26 **Fig. 2** and **Fig. S7a**). The MSA analysis of Os_cFUG1, ONIVA03G18050, and ORUFI03G17620 reveals two mismatches between Os_cFUG1 and ONIVA03G18050 (99.5% sequence identity), while Os_cFUG1 and ORUFI03G17620 are 100% identical (**Fig. S7b**). AtOTS1 and AtOTS2 are placed into cluster 9 and 10 and have no direct orthologous mapping to OsOTS1 and OsOTS2, in clusters 11 and 13 (**Fig. 2b & 2c**), indicating the independent duplication of OTSs in *Arabidopsis* and rice. OsOTS1 and OsOTS2 are in sister clades and share a common evolutionary origin. Within each OsOTS clade (clusters 11 and 13, **Fig. 2b & 2c**), there are orthologous ULP from RPRP rice (varieties of *Oryza sativa*), as well as other *Oryza* species, suggesting that the divergence of OTS1 and OTS2 occurred before speciation events within the *Oryza* genus. The divergence of OTS1 and OTS2 in monocot species probably pre-dated the *Oryza* genus, as there are orthologues of OTS2 in the nearest outgroup of the genus *Oryza*; cutgrass (LPERR01G25870.1; clusters 12 and LPERR12G15320.1; cluster 13, **Fig. 2b & 2c**), although a further exploration of the evolution of OTS genes is beyond the scope of this work.

### Sequence and structural analyses of ULPs in wild and domesticated rice

The catalytic domains of ULPs have crucial catalytic sites (HxDxC), which are important for deSUMOylation (Kurepa et al., 2003). We identified the protein domains in the representative ULP candidates (identified from pan-genes) using InterProScan and assessed the predicted disordered regions via metapredict (Emenecker et al., 2021). Disordered regions are indicated by lighter blue shades (values close to 1), and the more ordered regions are shown in darker blue (**Fig. 3a**). The ULP catalytic domains are typically located closer to the C-termini of the proteins (**Fig. 3b**). There are overlapping homologous superfamilies of other domains such as Papain-like cysteine peptidase superfamily (IPR038765), which are shown in light purple (**Fig. 3b**). These ordered regions align with the ULP1 protein domains as suggested by InterProScan, suggesting that there are potential functional domains in these ULPs at the C-termini as expected.

**Fig. 3.**
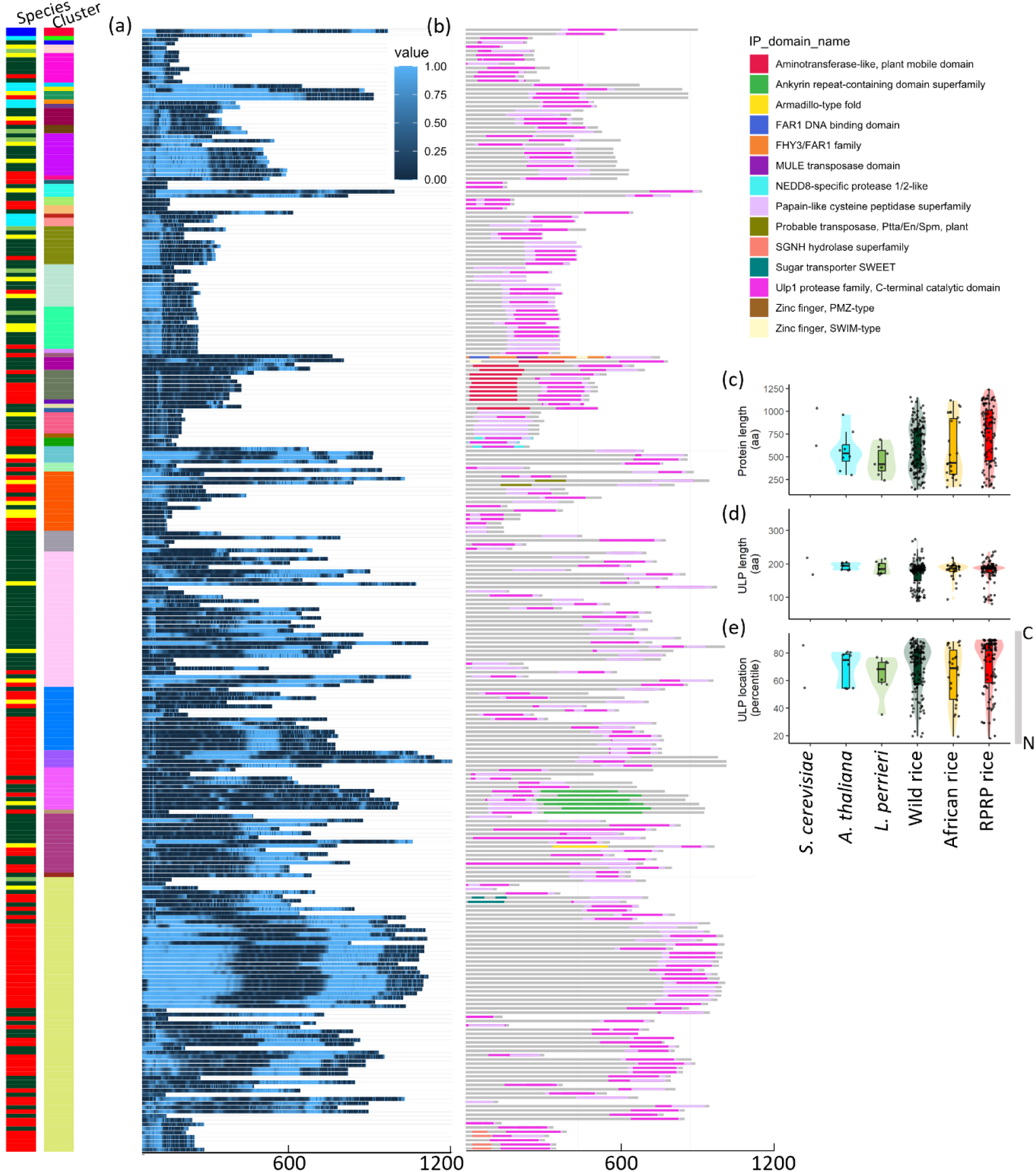
ULP protein domains architecture. **a**) Disorder regions prediction by metapredict. Scores are ranging from 0 to 1 (light blue) which indicates disorder regions. **b**) Protein domains identified by InterProScan. The ULP proteins are ordered by the gene tree in Fig. 3 .**c**) The length (amino acids) of ULPs. **d**) The length of ULP domains (amino acids). **e**) the ULP domains position on the ULP proteins. Colours in species annotation bar and Figs c and d represent species; *S. cerevisiae*, *A. thaliana, L. perrieri,* wild rice, African rice, and MAGIC16 rice (which includes Temperate Japonica; Temjap, Tropical Japonica;Tropjap, Aro rice, Aus rice, Indica rice).

The length of candidate ULPs varies across the species. RPRP genomes have relatively longer ULP protein sequences (**Fig. 3c**), also indicating larger protein structures than wild rice. The lengths of the ULP domains are similar amongst all species and varieties (**Fig. 3d**); however, the location of ULP domains within the proteins is variable (**Fig. 3e**). In yeast, there are two types of ULPs. The ScULP1 and ScULP2 have their domains located at C-terminus and in the middle of the protein, respectively. This feature seems to be conserved in *Arabidopsis* and rice as there are two populations of ULP locations on proteins (**Fig. 3e**). In rice, however, the majority of ULPs have their domains located at the C-terminus (**Fig. 3b & 3e**). The protein domain composition and disorder region predictions for all ULPs are presented in **Fig. S7**. We also display the chromosomal locations of all ULPs within the RPRP set in **Fig. S8**, demonstrating that most ULPs are consistently located on chromosomes across genomes, with some exceptions.

To assess whether the identified ULPs could potentially function as SUMO protease, we used Alphafold2 to predict the structures of the representative RPRP ULP representative proteins and all ULPs in wild rice. We measured the distances between the three residues that form the ULP catalytic site and compared them to the known ULP catalytic distances from (ScULP1, ScULP2, AtOTS1, AtOTS2, AtELS1, AtELS2, AtEDS4, AtFUG1, AtSPF1, AtSPF2, OsOTS1, and OsOTS2). Protein structures of RPRP ULPs that exhibited incorrect catalytic site conformations at the catalytic site, potentially impairing enzymatic activity or substrate binding, were excluded from the list (**Fig. S3**). The catalytic sites of ScULP1, AtOTS2, OsOTS2, and Os_nFUG1 are shown in **Fig. 4a-d**. The average catalytic distances (angstrom, Å) between H and D is 9.22, H and C is 6.39, and D and C is 7.91 (**Fig. 4E, Table S4**). The catalytic site distances in the pan-gene representatives (New) and wild rice (Wild) were similar and not significantly different from the known ULP reference distances (**Fig. 4e**), suggesting likely SUMO protease activity. However, the D to C distances showed slight variations from the references, suggesting some possible differences between wild and domesticated rice (**Fig. 4e**).

**Fig. 4.**
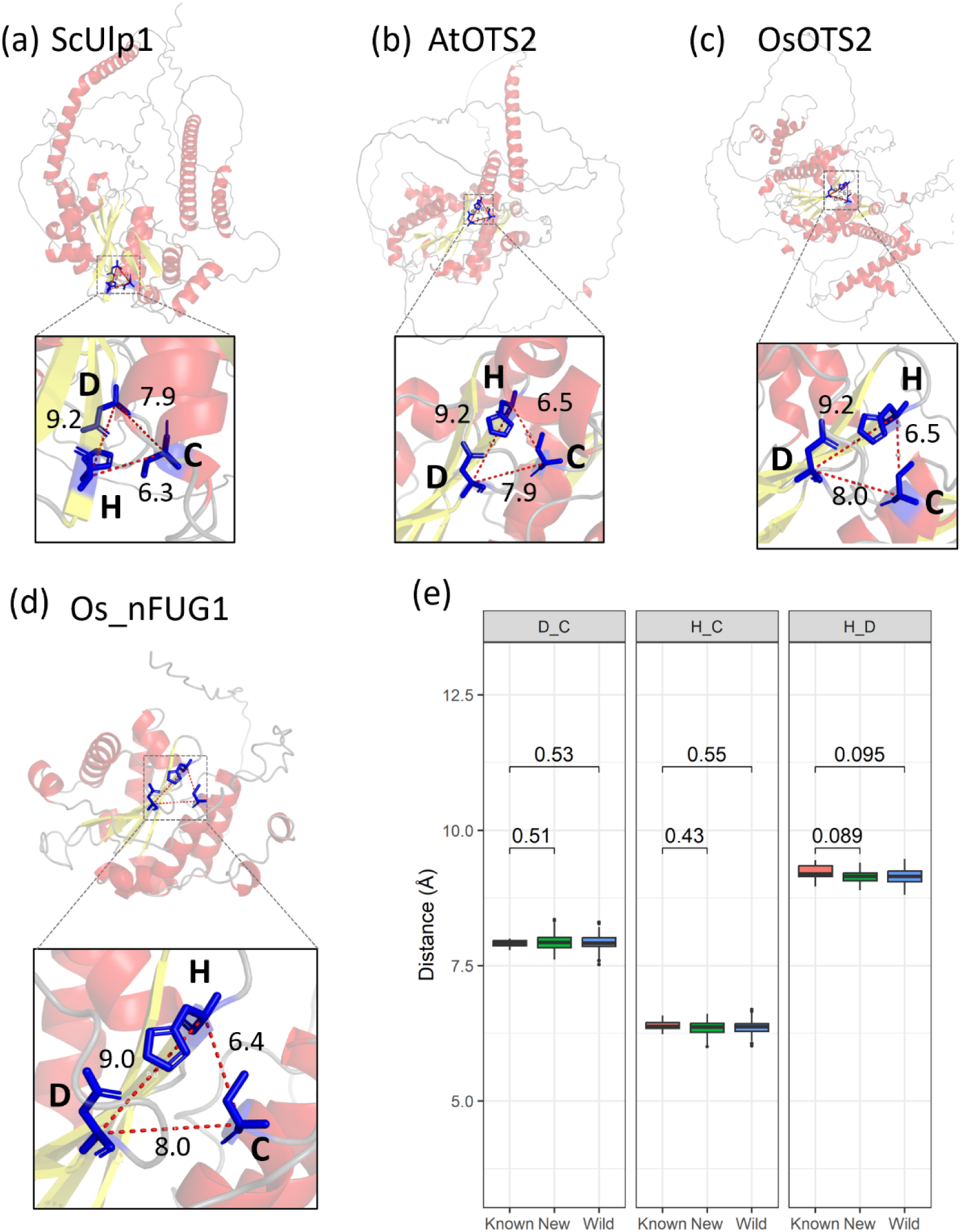
Protein structures predicted by ColabFold of **a)** ScULP1, **b)** AtOTS2, **c)** OsOTS2 and **d)** Os_nFUG1. **e)** The lengths (angstrom, Å) between the three catalytic sites (amino acids) of known ULPS (ScULP1, ScULP2, AtOTS1, AtOTS2, AtELS1, AtELS2, AtEDS4, AtFUG1, AtSPF1, AtSPF2, OsOTS1, OsOTS2). Sc: *S. cerevisiae*, At: *A. thaliana*, Os: *O. sativa* with novel ULPs in domesticated (New) and wild (Wild) rice. Statistical analysis was done with non-parametric Wilcoxson’s test and p-values are shown.

### Transcriptomics evidence of novel ULPs in O. sativa Japonica

To evaluate the expression of the identified ULPs, we examined their transcript levels using data from the plant RNAseq database (Yu et al., (2022): https://plantrnadb.com/ricerna/). We retrieved the FPKM expression values for the identified ULPs (RGAP models) from RNA-seq data across 631 samples of shoots, leaves, and roots under normal conditions (**Table S5**). These values were then scaled (z-score) for within-ULP comparison (**Fig. 5a**), and the z-scores for the three tissues were averaged (**Fig. 5b**). Hierarchical clustering reveals distinct tissue-specific expression patterns, highlighting the differential regulation of genes in the three rice tissues. Notably, several genes were predominantly expressed in the shoot, while *LOC_Os07g32090 (OsULP43c)*, *LOC_Os04g54670 (OsULP44a)*, *LOC_Os12g36580 (OsULP40d)*, and *LOC_Os09g12480 (OsULP45a)* exhibited relatively higher expression in the roots (**Fig. 5a-b**). To compare gene expression levels between ULPs, we scaled FPKM values across all the ULPs within the RNA-seq data. The ULPs that exhibited high expression across all three tissues under normal conditions include *LOC_Os12g01290 (OsULP36)*, *LOC_Os06g29310* (*OsOTS1*), *LOC_Os01g25370* (*OsELS1*), *LOC_Os05g11770* (*OsSPF1*), *LOC_Os03g22400* (*Os_cFUG1*), *LOC_Os11g01180 (OsULP37a)*, *LOC_Os11g10780* (*Os_nFUG1*), and *LOC_Os12g41380* (*OsOTS2*). In contrast, another known ULP, *LOC_Os03g29630* (*OsELS2*), displayed relatively lower expression, but its expression level still remained close to the mean (z-score = 0) (**Fig. S9a**).

**Fig. 5.**
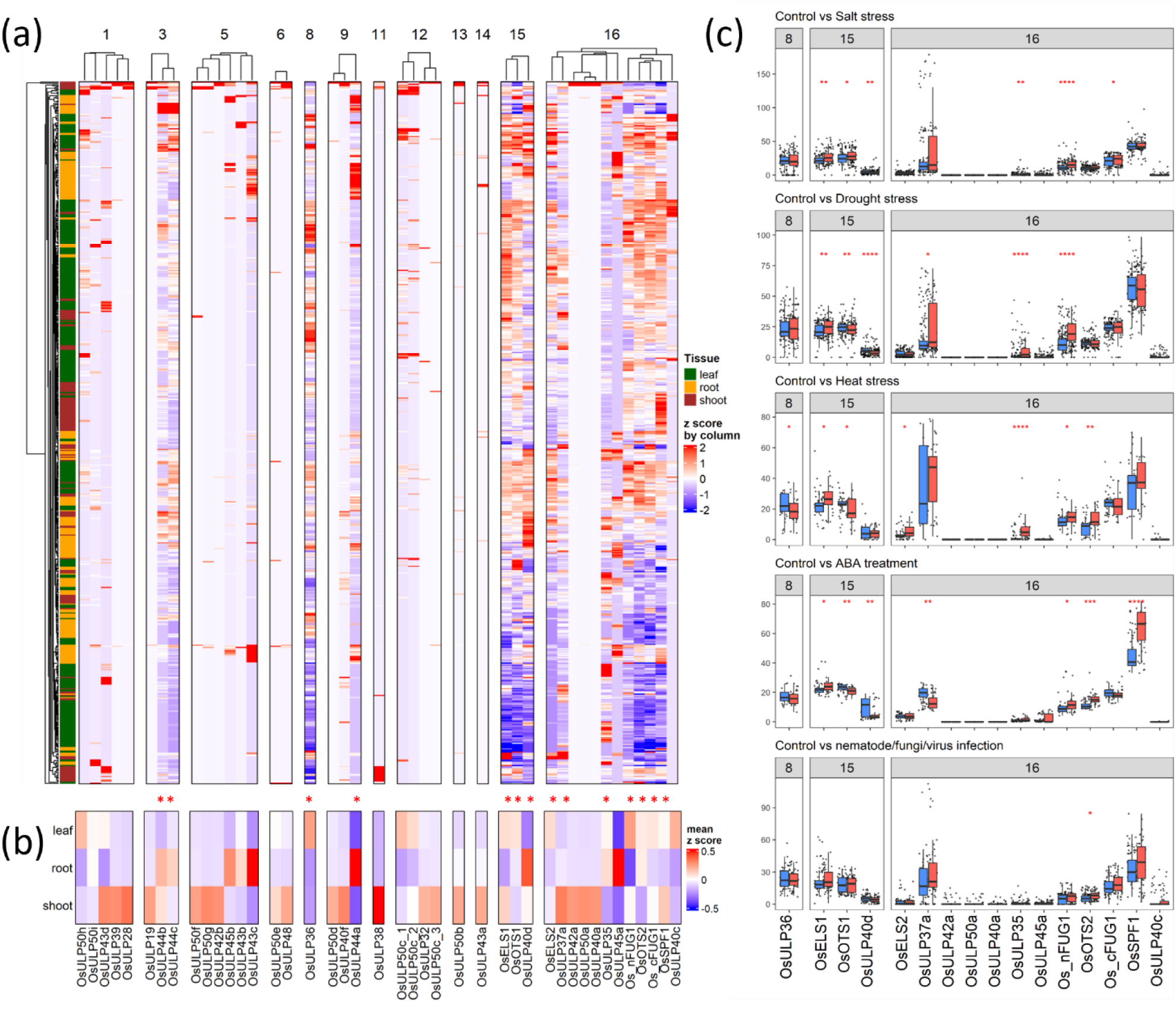
Transcriptomic evidence of the novel ULPs in *O. sativa* Japonica Nipponbare. **a)** A total 631 publicly available RNA-seq dataset of *O. sativa* leaves (315), shoots (116), and roots (200) showing tissue-specific patterns of the novel ULPs. The FPKM values were scaled by genes (z-score by column) then clustered by tissues (row) and within the ULP domain occupancy groups (1-16). **b)** Mean of z-score values of ULP expressions by tissues. The ULP genes that were expressed (FPKM >0) in over 50% of the samples are indicated by asterisk (*).**c)** The FPKM values of the 16 ULPs under control (blue) and 5 stress treatments (red): salt (148), drought (188), heat (54), nematode/fungi/virus infection (105), and ABA (45). See RNA-seq details in **Table S6-10**. The mean FPKM values between control and stress conditions were statistically compared using Wilcoxon’s test. Significance levels were indicated as follows: p ≤ 0.05 (*), p ≤0.01 (**), p ≤ 0.001 (***), and p ≤ 0.0001 (****). Only the conditions that have gene expression (FPKM > 0) in over 50% of the samples were included in the statistical test.

OTS1 and OTS2 have been shown to respond to salt and drought stress in both *Arabidopsis* and rice (Srivastava et al., 2017, 2018). Additionally, both OTS proteins play roles in bacterial pathogen responses in *Arabidopsis* (Bailey et al., 2016). To determine if other identified rice ULPs also respond to selected abiotic or biotic stresses, we assessed RNA-seq data from treatments including salt (148 samples), drought (188 samples), heat (54 samples), nematodes (*Meloidogyne graminicola*, *Hirschmanniella*), mycorrhizal fungi and Rice Stripe Virus (RSV) infection (105 samples), and ABA (45 samples), along with their respective controls (**Tables S6–S10, Fig. S9)**.

Overall, the highly expressed ULPs demonstrated responsiveness to the abiotic stresses (**Fig. 5c**). Under salt conditions, we observed significant up-regulation (Wilcoxon’s test, p-value < 0.05) of *LOC_Os01g25370* (*OsELS1*), *LOC_Os06g29310* (*OsOTS1*), *LOC_Os11g10780* (*Os_nFUG1*), and *LOC_Os03g22400* (*Os_cFUG1*). Under drought conditions, *LOC_Os01g25370* (*OsELS1*) and *LOC_Os11g10780* (*Os_nFUG1*) were up-regulated, similar to their response under salt treatment. In contrast, *LOC_Os06g29310* (*OsOTS1*) was significantly down-regulated. Heat stress significantly up-regulated *LOC_Os01g25370* (*OsELS1*), *LOC_Os03g29630 (OsELS2)*, *LOC_Os10g33450 (OsULP35)*, *LOC_Os11g10780* (*Os_nFUG1*), and *LOC_Os12g41380* (*OsOTS2*). On the other hand, heat stress caused the significant down-regulation of *LOC_Os12g01290 (OsULP36) and LOC_Os06g29310* (*OsOTS1*). Under ABA treatments, a central phytohormone involved in responding to abiotic stresses, we observed significant upregulation of *LOC_Os01g25370* (*OsELS1*), *LOC_Os11g10780* (*Os_nFUG1*), *LOC_Os12g41380* (*OsOTS2*), and *LOC_Os05g11770* (*OsSPF1*). In contrast, *LOC_Os06g29310* (*OsOTS1*) and *LOC_Os11g01180 (OsULP37a)* were down-regulated.

Under biotic stress condition, only OsOTS2 showed significantly up-regulation under the nematode/fungi/virus infection (**Fig. 5c, Table S6-S10**). In addition, three ULPs (*LOC_Os10g06910*, *LOC_Os09g08440*, and *LOC_Os02g27280*), which are ULP-domain conserved across RPRP pan-genome but exhibit low or zero expression levels in the other stresses, showed slightly increased transcription under the pathogen infection. This suggests that these ULPs may be involved in biotic stress responses.

## Discussion

ULPs are one of the key components in SUMOylation process because of their dual functions in maturing SUMO proteins and cleaving SUMOylated proteins (Yates et al., 2016). In *O. sativa* Japonica Nipponbare, the identification of ULPs has been previously reported, with some speculation of the ULP expansion during domestication (Yates et al., 2016; Srivastava et al., 2016; Garrido et al., 2018). However, previous methods used relatively simple sequence-based analysis to determine the count of ULP genes/proteins. In this study, we greatly expanded on previous research by taking a multi-pronged approach, including sequence searching, domain identification, phylogenetics and structural bioinformatics to validate the presence of the specific protease active site. We explored the ULP gene/protein set within the IRGSP RefSeq, and within a population-aware pan-genome set (RPRP), demonstrating that the family expansion (compared to model organisms) is consistently observed in multiple varieties. We also observed within the pan-genome analysis that ULP proteins are commonly found with single or low occupancy (cloud genes) across the pan-genome, which tends to be indicative of genes under selective breeding pressure. The pan-gene set analysis of the RPRP genomes also identified particular protein domains as invariable, partially variable and highly variable across the pan-genome (based on the mean domain occupancy being <10, across 16 genomes) – with the highly variable domains characterised by those involved in biotic stress responses (like NB-ARC and LRR) (Contreras-Moreira et al., 2025). The ULP protease domain falls into the highly variable domain, as the mean domain occupancy is 5.94, with an enrichment of single copy genes (present in only one genome).

The count of ULP genes in cultivated rice is higher than wild rice. The rice IRGSP RefSeq contains 45 ULPs, and the RPRP genomes (representing the genetic diversity of Asian rice) have ULP counts ranging between 32-39. The higher count in the IRGSP RefSeq might be due the presence of a larger number of genes called from the three sets (RAP-DB, RGAP and Gramene), compared to a single gene model set in the RPRP genomes.

In contrast, wild rice species have a broader range with ULP numbers ranging from 9-36 ULPs. *O. puntata* is an African wild rice species, considered as a weed (Wirjahardja et al. (1983), and contains the highest number of ULP (36) among wild rice. *O. puntata* is classified as BB genome, which is distantly related to RPRP rice (AA genome type). This indicates a divergence of ULP expansion in *Oryza* relatives. Interestingly, *O. punctata* is regarded as having pathogen-resistant traits and hence has been explored for breeding with cultivated rice varieties (Jena et al., 2016).

Asian rice was cultivated about 9000 years ago between two wild rice species: *O. rufipogon* and *O. nivara* (Fornasiero et al., 2022). Over time, beneficial traits such as grain quality, stress and pathogen resistance were selected through the breeding process (Hernández-Soto et al., 2021). Genes inherited from these wild ancestors have been identified as domestication-associated genes (Jing et al., 2023). *Os_cFUG1* is one of the domesticated genes derived from *O. rufipogon* (Jing et al., 2023). However, our phylogenetic analysis of the ULPs suggests that Os_cFUG1 is more closely related to ELS-type and the protein sequence is identical to the *O. nivara* orthologue. Based on this observation, we propose a novel *OsFUG1* (*Os11g0214001*, *LOC_Os11g10780*) and *OsELS3* (renaming the current *OsFUG1*; *Os03g0344300*, *LOC_Os03g22400*). In addition, the SUMO-conjugating enzyme OsSCE1a was identified as a key regulatory protein in 9YJ30 rice breeding (Yuan et al., 2025). The 9YJ30-F_2_ hybrid was obtained from backcrossing between F_1_ (a hybrid from a wild rice *O. rufipogon,* Yuanjiang rice, and an elite *O. sativa* indica, 93–11 line) with 93–11 (parent). OsSCE1a has been shown to associate with functional “stay-green” traits and growth duration in this 9YJ30-F_2_ rice (Yuan et al., 2025). These findings highlight the importance of SUMOylation processes and their likely selection during rice domestication.

The public transcriptomic data analysis revealed gene expression patterns of the identified ULP in three main tissues of rice: leaves, shoots and roots (**Fig. 5**). The highly expressed ULPs have domain occupancy generally in at least 15 out of 16 of the pan-genome set. However, another highly expressed ULP, LOC_Os12g01290, with 8 domain occupancy, is found in Tempjap, Tropjap, Aro, and Aus rice varieties but not in Indica varieties except *Os*Lima. This gene, however, has an unclear gene model, making it uncertain whether any of the source models are accurate (**Fig. S10**). Given the evidence of transcription, further investigation, including gene cloning, protein expression and SUMO protease assays, will be necessary.

SUMOylation modifies proteins, e.g. transcription factors, to regulate their functions (promote or inhibit) during stress responses (Han et al,. 2017). For example, SUMOylation of the rice DELLA protein SLR1 at lysine 2 has been found to modulate transcriptional responses and improve yield under salt stress (Fernandes et al., 2024). SUMO proteases, on the other hand, works oppositely, by deSUMOylating the target proteins (e.g. transcription factors) and resetting the system.

Five ULP genes from our list: *Os03g0365250 (LOC_Os03g24960)*, *Os03g0365200 (LOC_Os03g24990)*, *Os04g0583400 (LOC_Os04g49394), Os10g0474000 (LOC_Os10g33450)*, and *Os10g0388600 (LOC_Os10g24954)* (**Table 1**) were identified as salt-related candidate genes from comprehensively integrating quantitative trait loci (QTL) and transcriptomic analyses of salt treatments in *O. sativa* whole seedings, excluding root samples (Phosuwan et al., 2024), suggesting functions of these proteases under salt stress. However, their transcription levels were low in our list of RNA-seq datasets (**Fig. S9b**), possibly due to the temporal induction of gene expression. In contrast, the two known salt-responsive proteins OsOTS1 (Os06g0487900) and OsOTS2 (Os12g0606900) were not included in their list because these proteins are expressed and responsive to salt exclusively in roots (Srivastava et al., 2017). This highlights the tissue specificity of ULPs in *O. sativa*.

OTS1 and OTS2 act as negative regulators of salicylic acid (SA) signalling in *Arabidopsis* when challenged with the bacterial pathogen *Pseudomonas syringae* (Bailey et al., 2016). During *P. syringae* infection, SA accumulation induces the degradation of OTS1 and OTS2 proteins. Once the infection is managed, the levels of OTS1 and OTS2 proteins are restored to regulate the expression of *ISOCHORISMATE SYNTHASE1* (*ICS1*) to prevent excessive responses to the pathogen (Bailey et al., 2015). We also observed significant upregulation of *OsOTS2* transcripts but non-significant upregulation of *OsOTS1* in rice infected with nematodes, fungi, and viruses (**Fig. 5c**). This suggests that *OsOTS2* may be upregulated to restore protein levels after overcoming the infection. However, we observed upregulation of other ULPs (**Fig. 5c**), which, although not statistically significant, may still play a role in the pathogen defence mechanism.

In conclusion, our analysis identified 45 ULPs in *O. sativa*. We also characterised ULPs in wild and RPRP rice at the pan-genomic level, providing evidence of potential functional ULPs and their transcript expression. Additionally, our study offers insights into the role of these ULPs in response to abiotic and biotic stresses. Experimentally characterising the deSUMOylation function of these novel ULPs using a SUMO protease assay (Srivastava, Zhang, & Sadanandom, 2016) and identifying their target proteins will be crucial for understanding their specificity. Finally, the novel ULPs identified in this study could serve as valuable targets for future rice breeding and other crop improvement.

## Supporting information

https://github.com/Kawinnat/ULP_identification_rice/blob/main/Supplementary_tables.xlsx

## Acknowledgements

KS’s PhD was supported by the DPST (Development and Promotion of Science and Technology Talents Project) scholarship from the Ministry of Higher Education, Science, Research and Innovation, Thailand. We would also like to acknowledge funding from BBSRC for the “SUMOCode” project [BB/V003534/1] and the “PanOryza” project [BB/T015691/1].

## Author contributions

KS, AS, RB, ES, CZ and AJ contributed to the conceptualisation, methodology, and formal analysis of the study. KS performed the analyses, created the visualisations, and wrote the original draft. KS, CZ, AS, RB, ES and AJ assisted in reviewing and editing the manuscript and gave final approval for publication.

## Data availability

The scripts used in this project are available on GitHub (https://github.com/Kawinnat/ULP_identification_rice).

## Supplementary Fig

**Supplementary Fig. 1.**
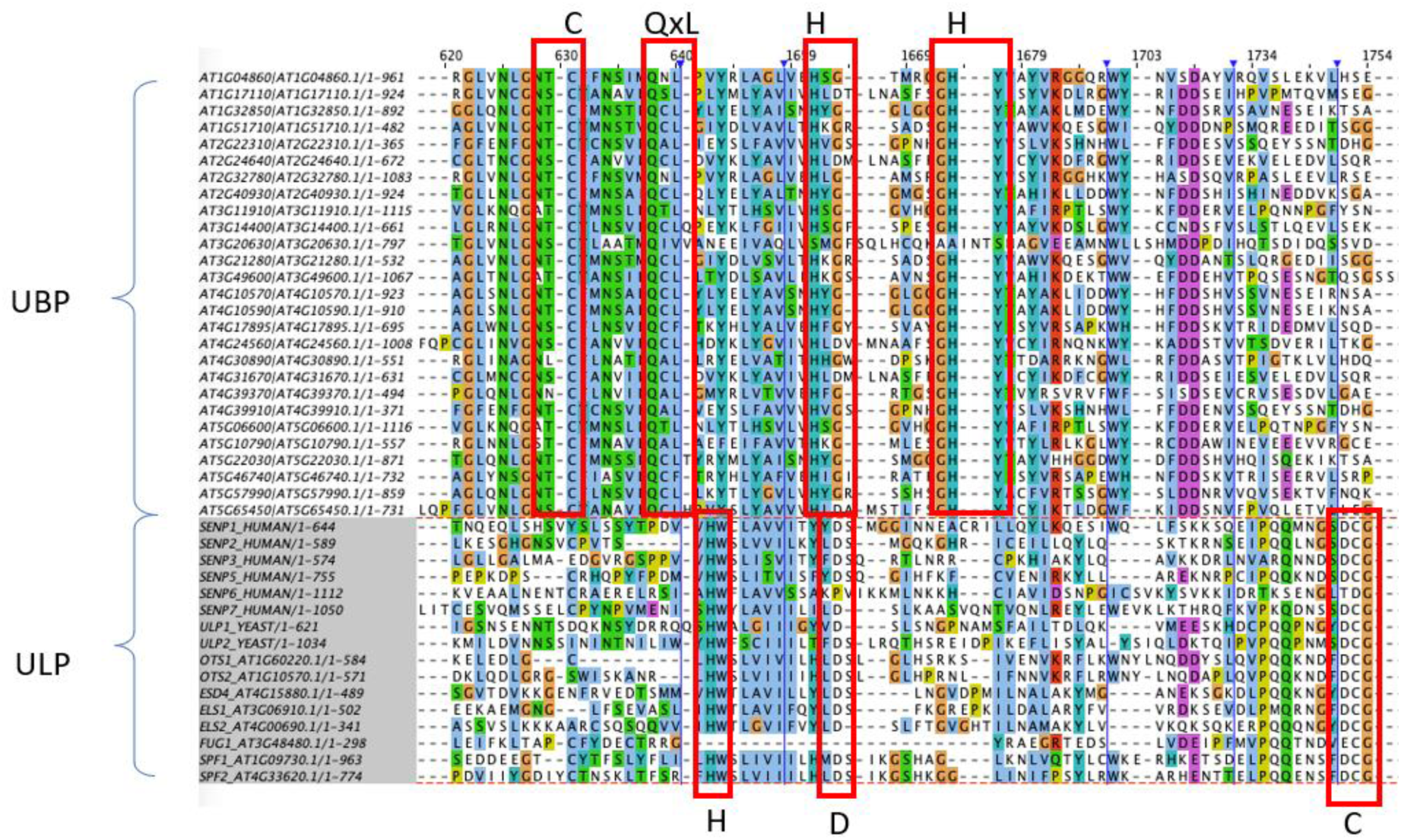
Multiple sequence alignment shows the catalytic sites of well characterized ULPs (SUMOylation) that distinguished from UBPs (Ubiquitination) in *A. thaliana*. The protein sequences of UBPs and ULPs were aligned (MUSCLE) and visualised in Jalview. The catalytic sites of ULPs and UBPs are highted in red. ULPs have HxDxC while UBPs have CxQxLxH forming catalytic sites (Zhou et al., 2017).

**Supplementary Fig. 2.**
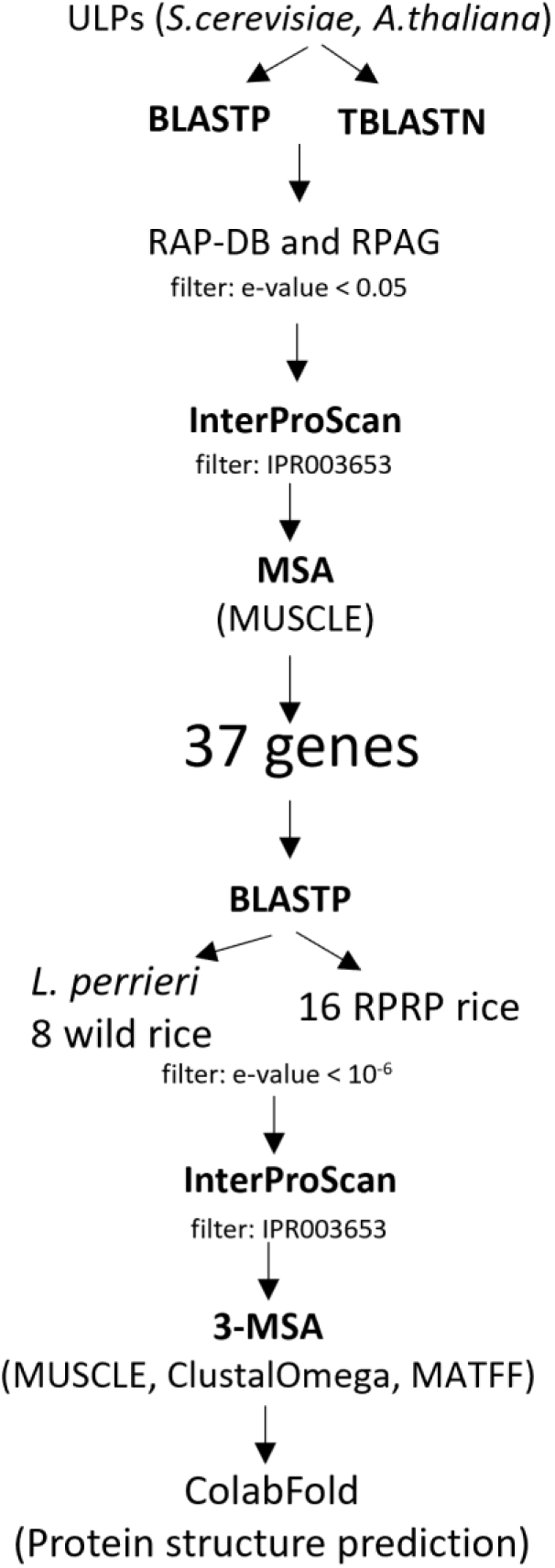
ULP identification workflow. Identification of ULPs by sequence-base and validation by structure-base searches in 2 rice reference genomes. Known ULP sequences from yeast and *Arabidopsis* were queried to BLASTP and TBLASTN searches in two *O. sativa* reference genomes (RAP-DB and RGAP). The significant sequence hits (e-value < 0.05) from BLAST results were scanned by InterProScan and only the sequences with ULP domains were retained. The ULP domain containing proteins were also obtained from the annotations RAP-DB and RGAP. The ULP candidates from sequence-base searches were aligned to known ULPs from yeast and *Arabidopsis* using MUSCLE and only the sequences with the right catalytic sites were carried on. We then searched for the ULPs orthologs in RPRP genomes and wild rice populations, using similar steps and cut offs; BLASTP and InterProScan. The proteins with ULPs catalytic domains were aligned with known ULPs from yeast and *Arabidopsis*. Lastly, the representative ULPs from pan-gene clusters and all identified ones in wild rice had protein 3D structures predicted, and the ones with correct catalytic areas were retained.

**Supplementary Fig. 3.**
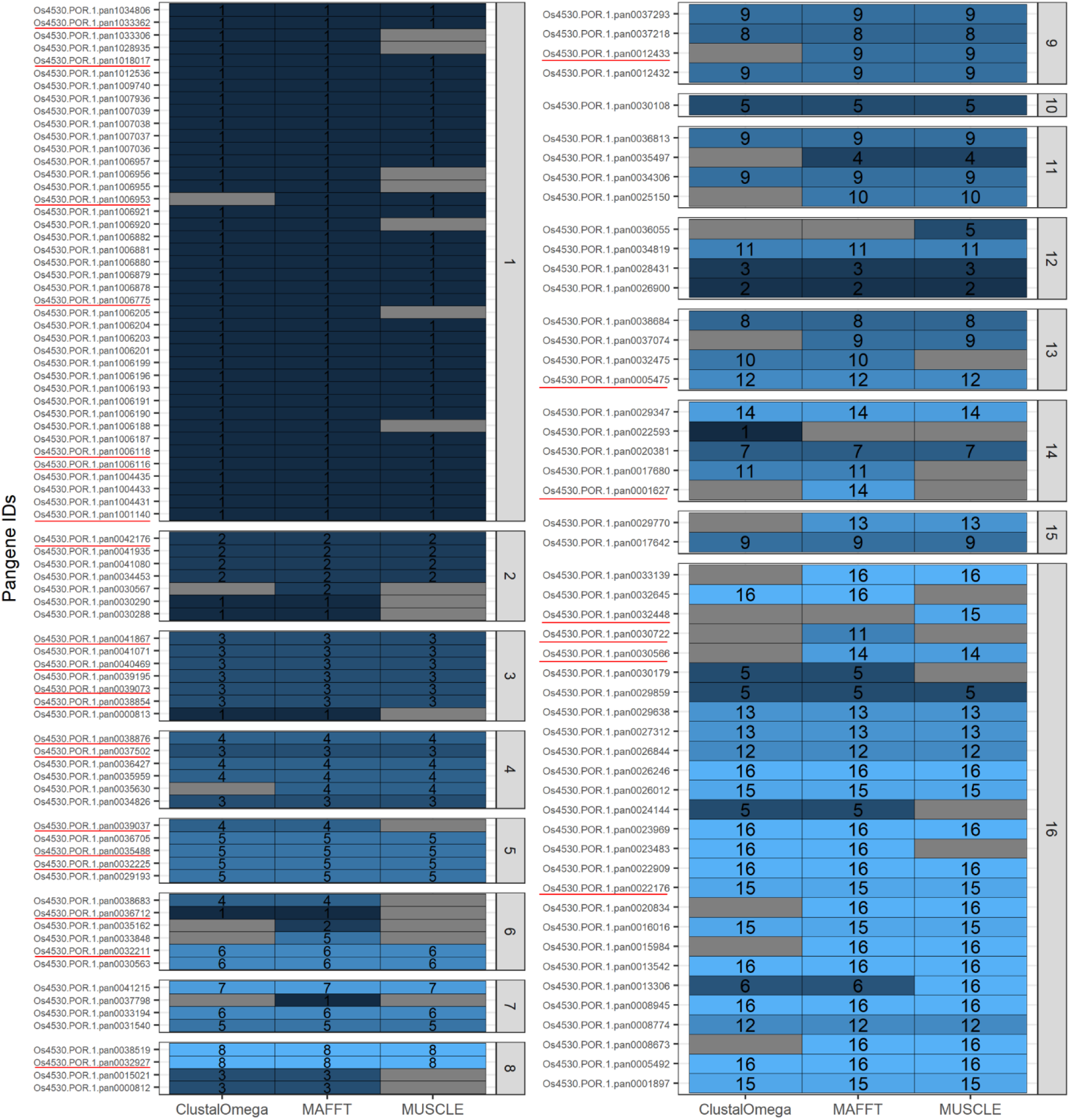
131 Pan-gene clusters of genes that harbour ULP domains and catalytic sites. The individual clusters were aligned by using three MSA tools (MUSCLE, MAFFT, and ClustalOmega) with known ULPs of yeast and *Arabidopsis*. The total number of genes that have correct catalytic sites are shown are also illustrated by the gradient of blues. The pan-gene clusters are separated by the “genome occupancy” within the RPRP populations. The pan-gene clusters for which their protein structure (AlphaFold) suggests wrong catalytic sites are underlined.

**Supplementary Fig. 4.**
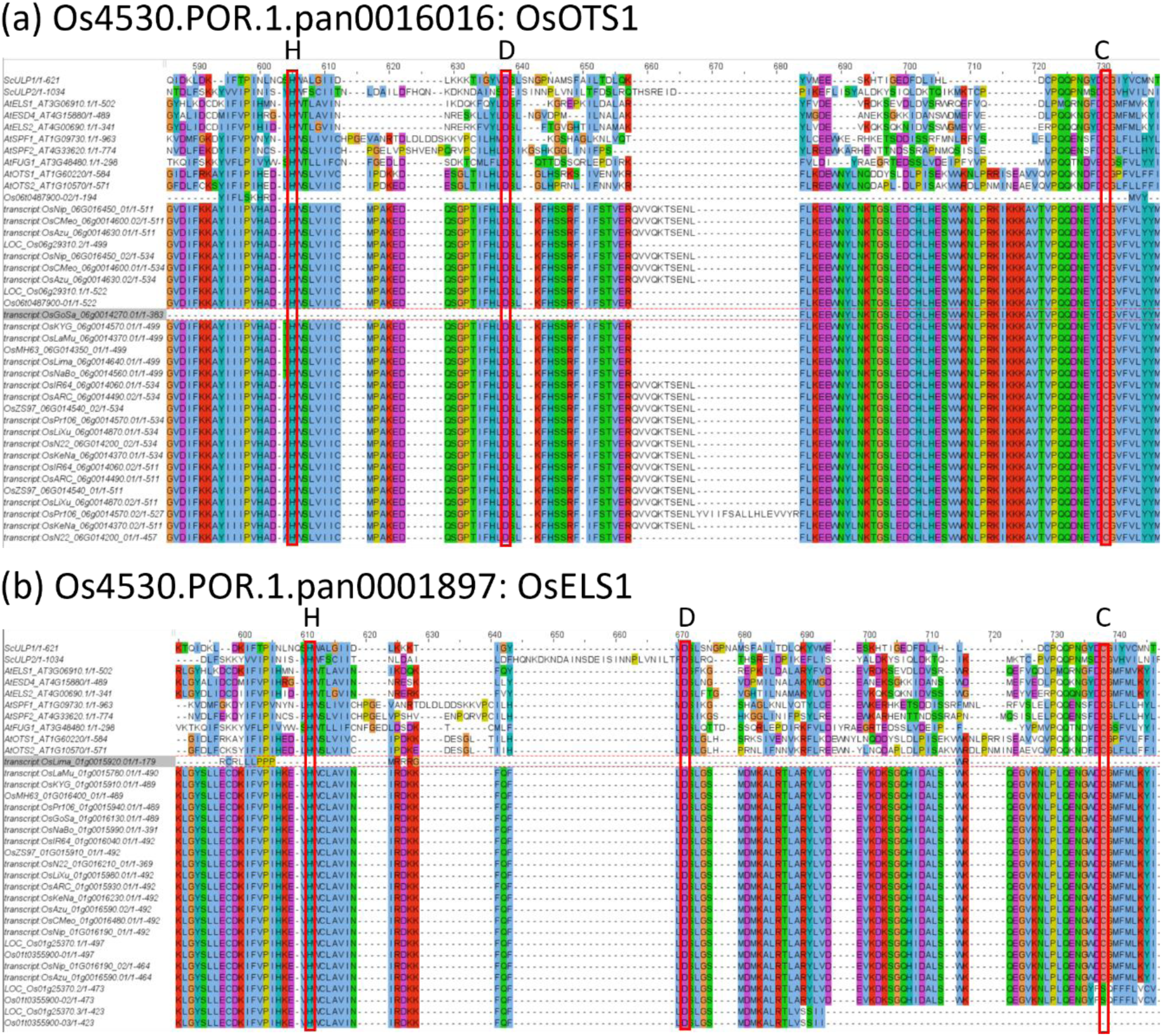
Multiple sequence alignments of a) OsOTS1 and b) OsELS1 pan-genes. The protein sequences were aligned using MUSCLE v5 and visualised in Jalview. The catalytic sites are indicated by red boxes.

**Supplementary Fig. 5.**
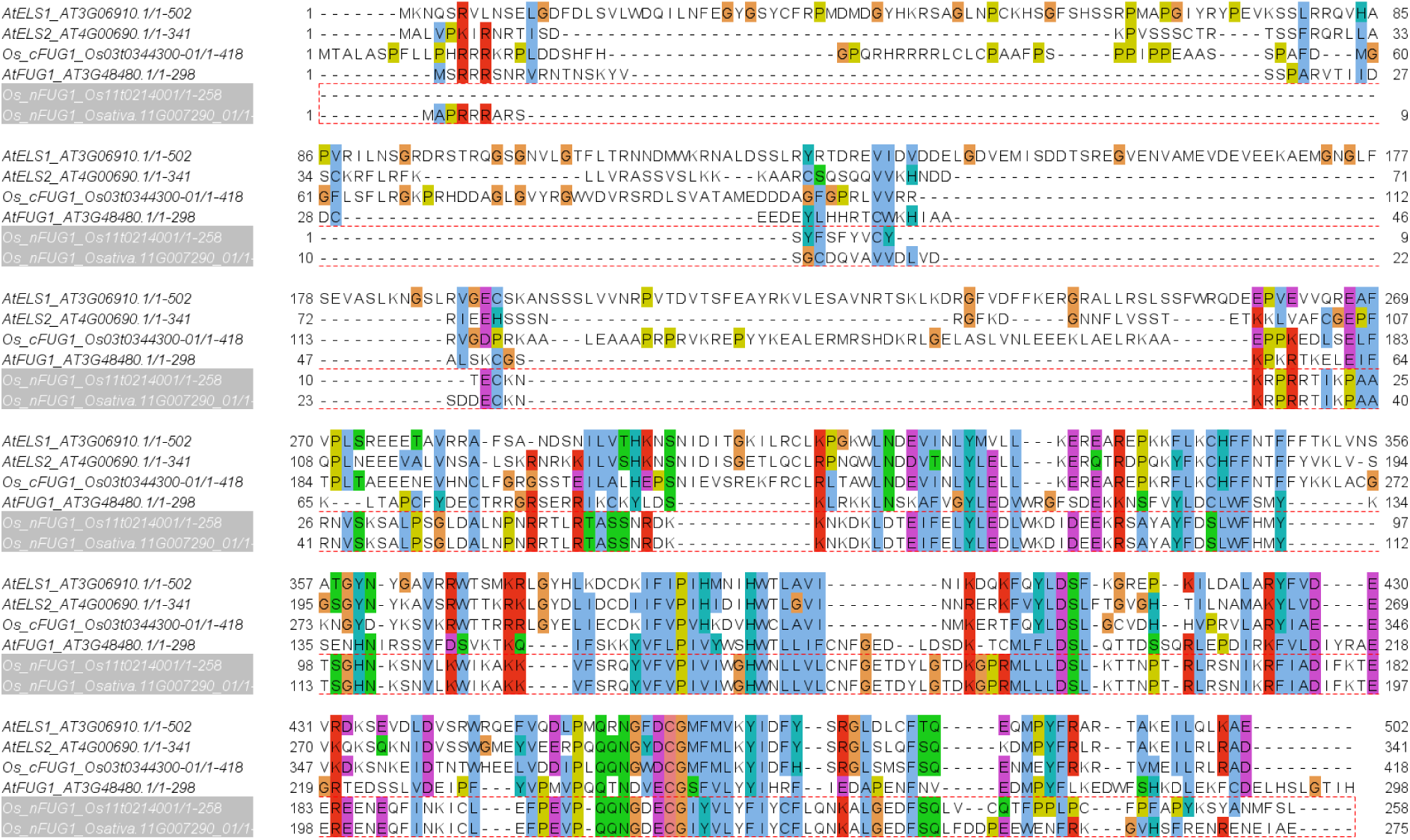
MSA of AtELS1/2, Os_cFUG1, AtFUG1, and Os_nFUG1 (RAP-DB and Gramene). The protein sequences were aligned using MUSCLE v5 and visualised in Jalview.

**Supplementary Fig. 6.**
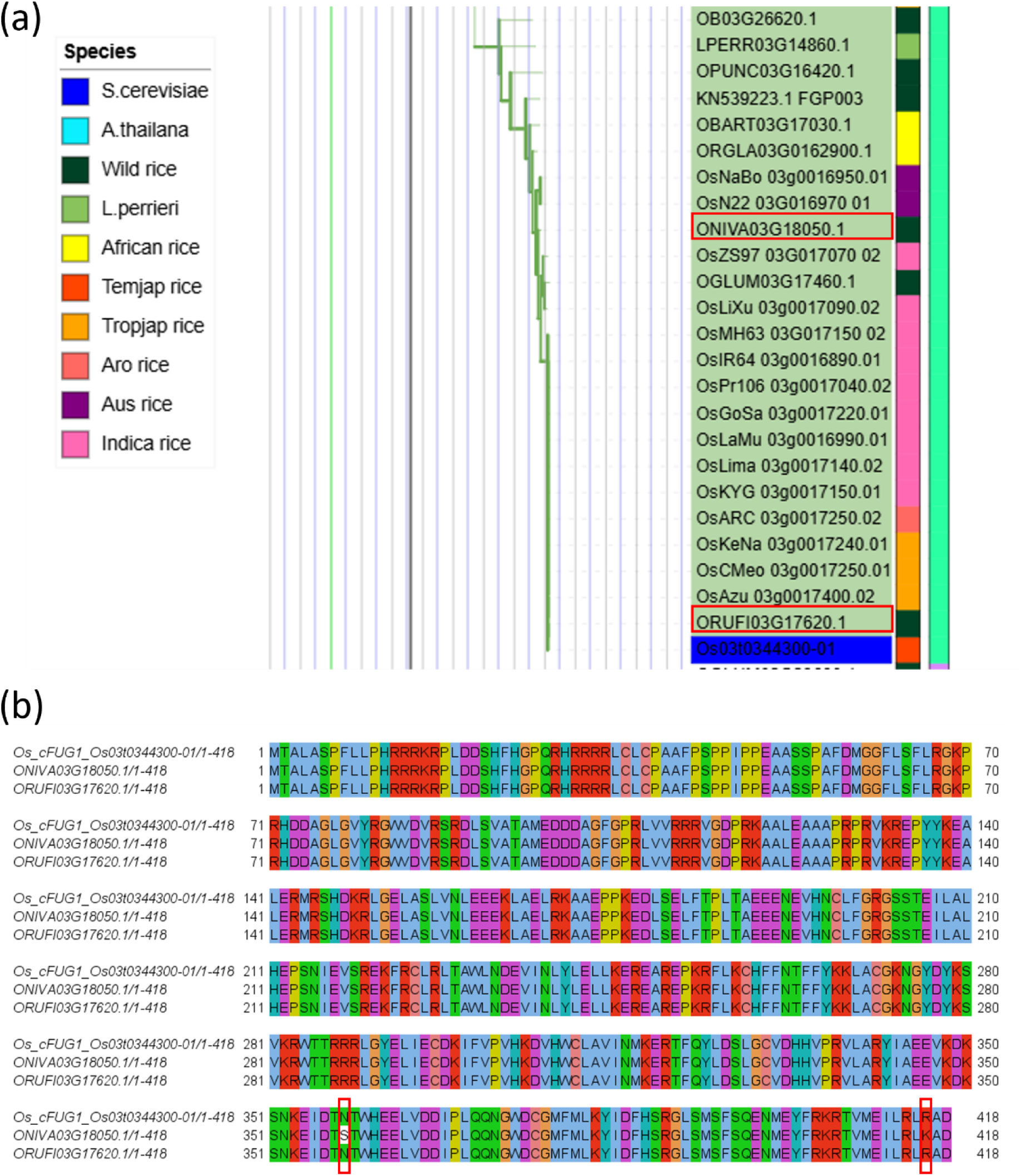
a) Cluster 26 showing the cluster of Os_cFUG1 b) Os_cFUG1 protein sequence alignment with its orthologues in *O.nivara* and *O.rufipogon*. The protein sequences were aligned with MUSCLE and two mismatches were highlighted in red boxes.

**Supplementary Fig. 7.**
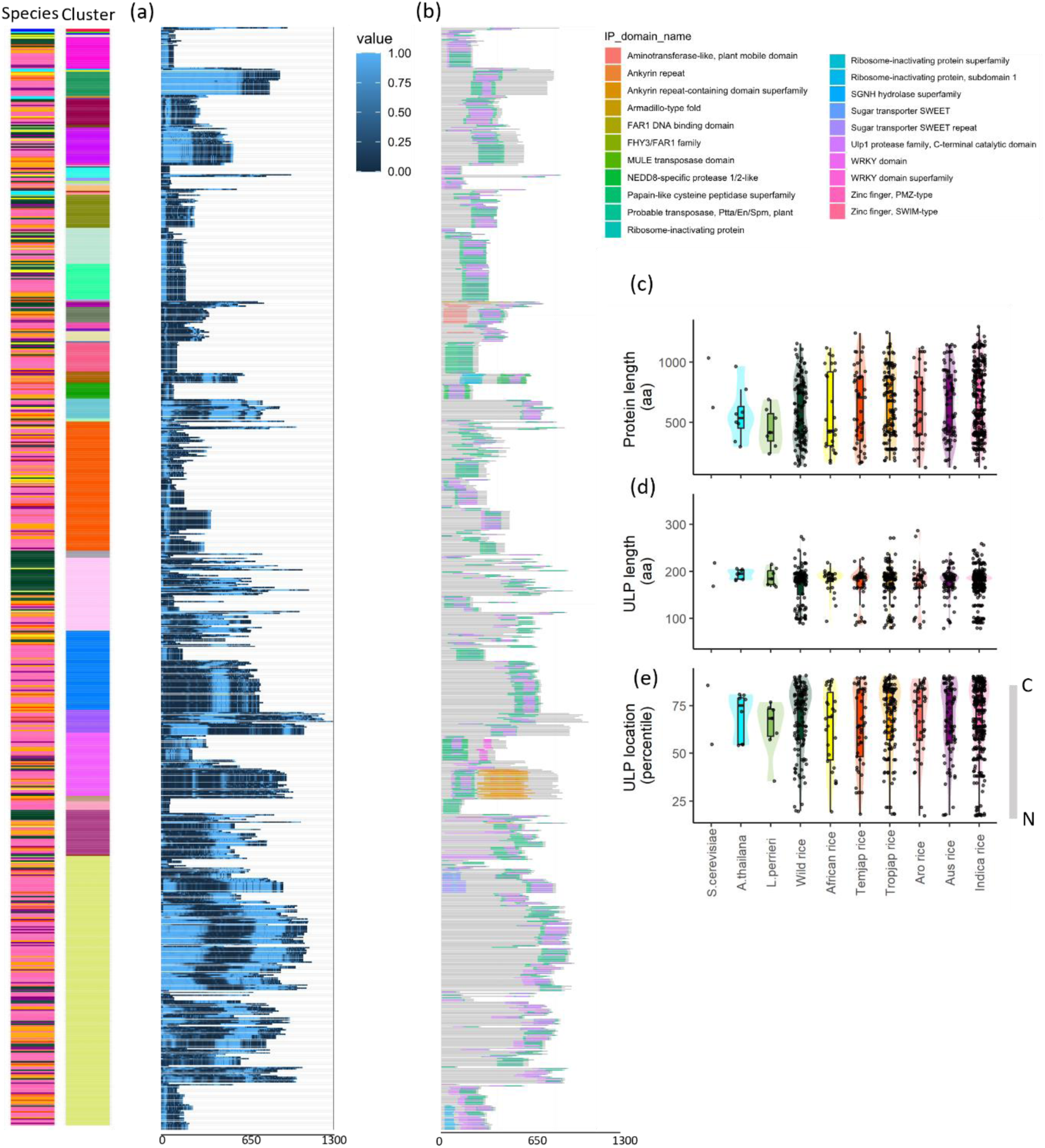
All ULP protein domains architecture. a) Disorder regions prediction by metapredict. Scores are ranging from 0 to 1 (light blue) which indicates disorder regions. b) Protein domains identified by InterProScan. The ULP proteins are ordered by the gene tree in Fig. 3 .c) The length (amino acids) of ULPs. d) The length of ULP domains (amino acids). E) the ULP domains position on the ULP proteins. Colours represent species; *S. cerevisiae*, *A. thaliana, L. perrieri,* wild rice, African rice, Temperate Japonica (Temjap), Tropical Japonica (Tropjap), Aro rice, Aus rice and Indica rice.

**Supplementary Fig. 8.**
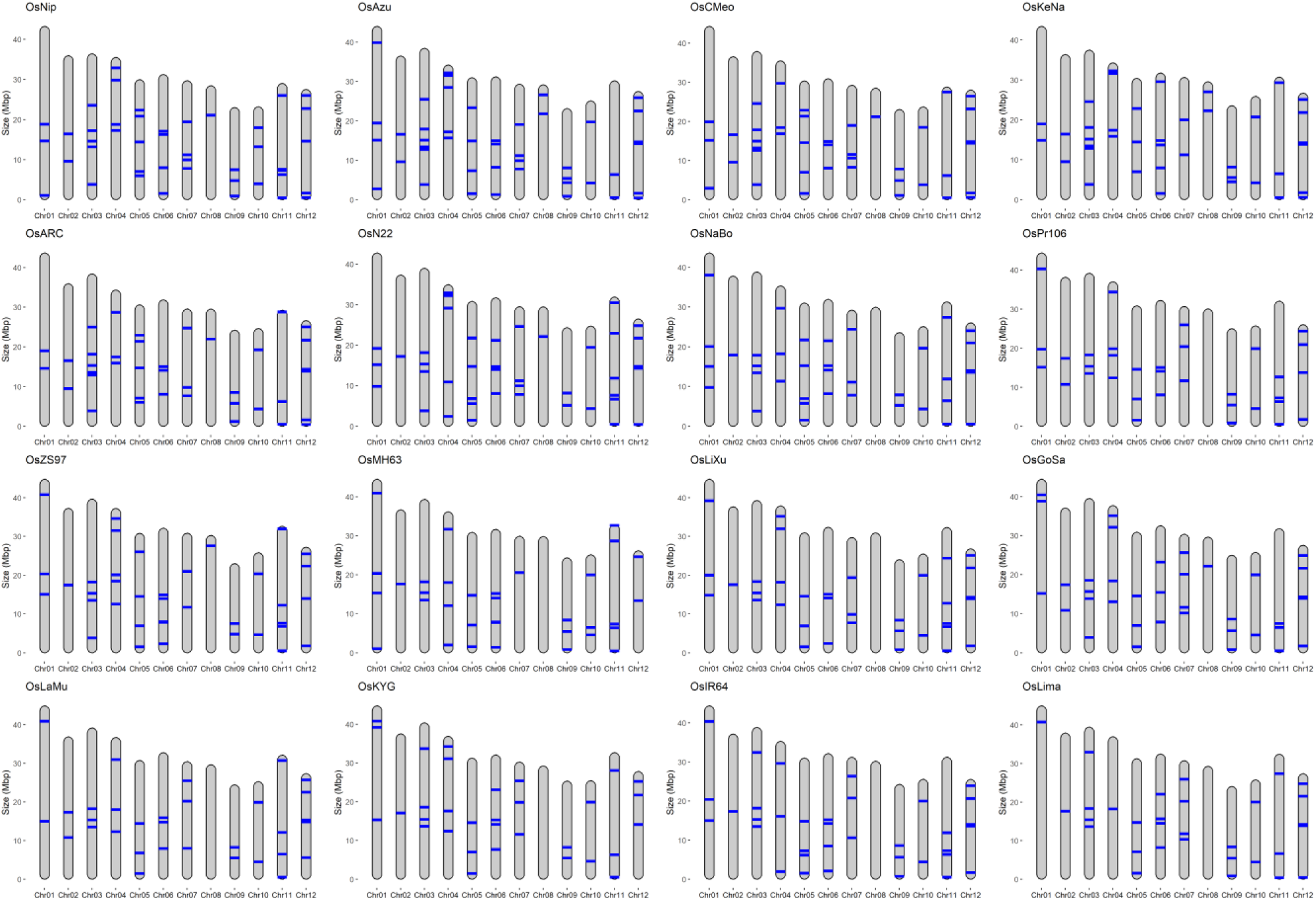
Genomic location of ULPs in RPRP rice population.

**Supplementary Fig. 9.**
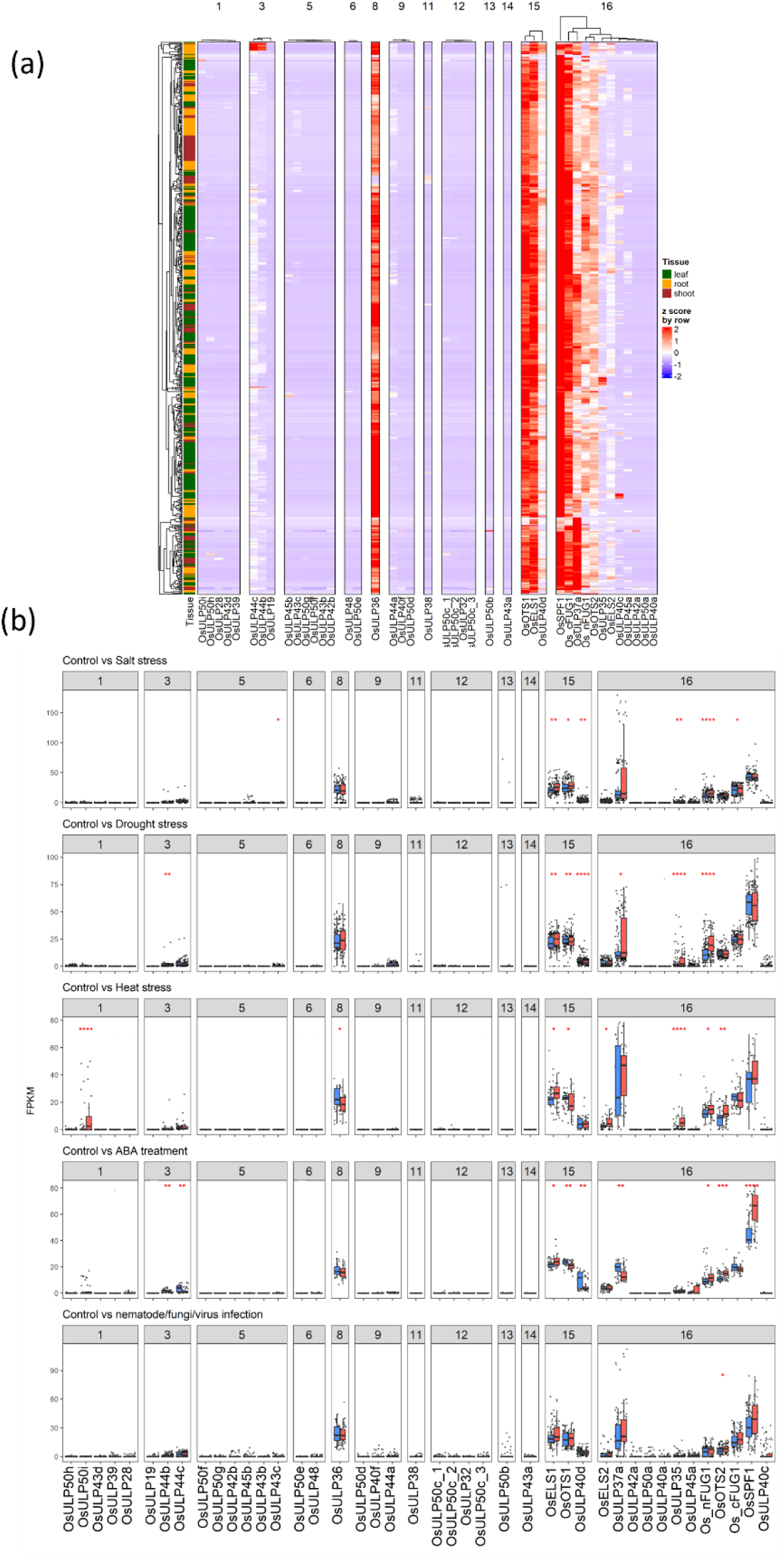
Transcriptomic evidence of the novel ULPs in *O. sativa* Japonica Nipponbare. a) A total 631 publicly available RNA-seq datasets of *O. sativa* leaves (315), shoots (116), and roots (200) showing tissue-specific patterns of the novel ULPs. The FPKM values were scaled by RNA-seq (z-score by row) then clustered by tissues (row) and within the domain occupancy groups (1-16). b) The FPKM values of all ULPs under control (blue) and 5 stress treatments (red): salt (148), drought (188), heat (54), ABA (45), and nematode/fungi/virus infection (105). See RNA-seq details in **Table S6-10**. The mean FPKM values between control and stress conditions were statistically compared using Wilcoxon’s test. Significance levels were indicated as follows: p ≤ 0.05 (*), p ≤0.01 (**), p ≤ 0.001 (***), and p ≤ 0.0001 (****). Only the conditions that have gene expression (FPKM > 0) in over 50% of the samples were included in the statistical test.

**Supplementary Fig. 10.**
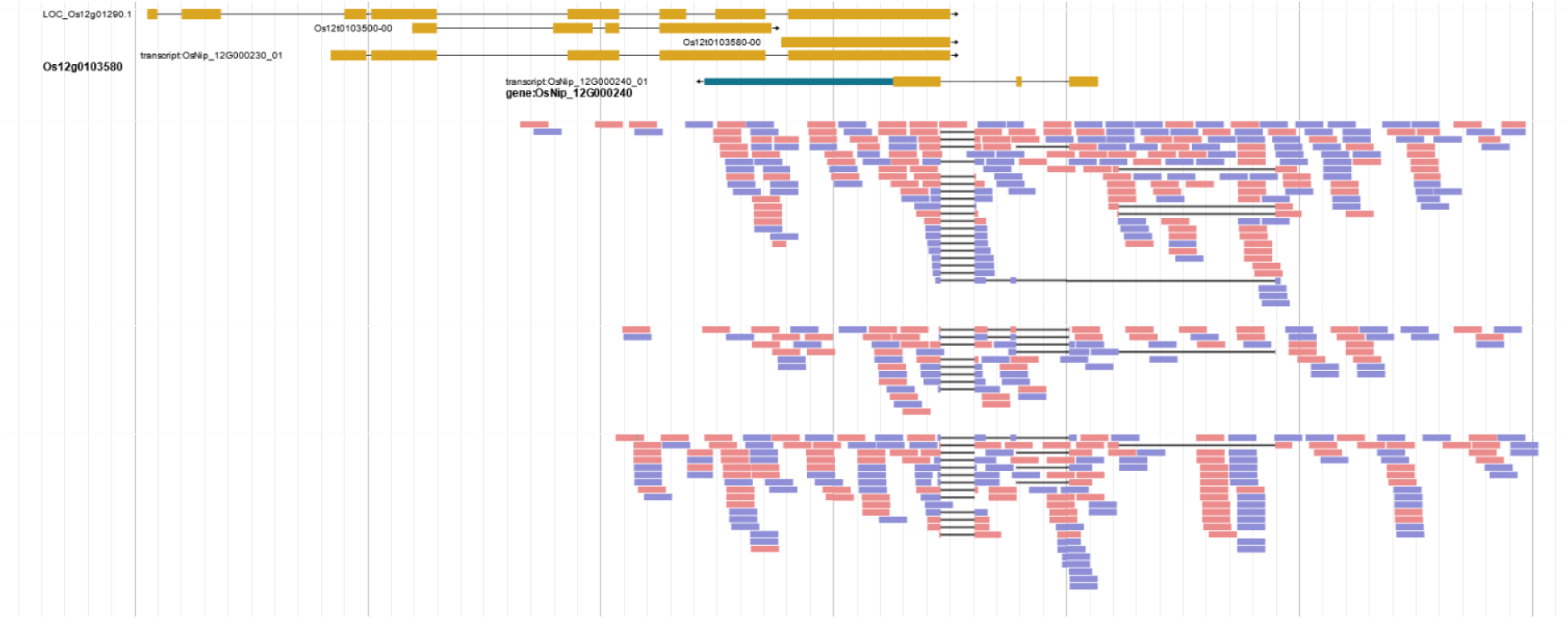
Screenshot of *LOC_Os12g01220* gene models derived from (https://panoryza.org/jbrowse/). The transcriptomic evidence is shown in pink and blue (pair-ended RNAseq).

## References

Augustine, R. C., & Vierstra, R. D. (2018). SUMOylation: re-wiring the plant nucleus during stress and development. Curr Opin Plant Biol, 45(Pt A), 143–154. 10.1016/j.pbi.2018.06.006

Augustine, R. C., York, S. L., Rytz, T. C., & Vierstra, R. D. (2016). Defining the SUMO System in Maize: SUMOylation Is Up-Regulated during Endosperm Development and Rapidly Induced by Stress. Plant Physiol, 171(3), 2191–2210. 10.1104/pp.16.00353

Bailey, M., Srivastava, A., Conti, L., Nelis, S., Zhang, C., Florance, H., Love, A., Milner, J., Napier, R., Grant, M., & Sadanandom, A. (2016). Stability of small ubiquitin-like modifier (SUMO) proteases OVERLY TOLERANT TO SALT1 and −2 modulates salicylic acid signalling and SUMO1/2 conjugation in Arabidopsis thaliana. J Exp Bot, 67(1), 353–363. 10.1093/jxb/erv468

Benlloch, R., & Lois, L. M. (2018). Sumoylation in plants: mechanistic insights and its role in drought stress. J Exp Bot, 69(19), 4539–4554. 10.1093/jxb/ery233

Blackburne, B. P., & Whelan, S. (2013). Class of multiple sequence alignment algorithm affects genomic analysis. Mol Biol Evol, 30(3), 642–653. 10.1093/molbev/mss256

Camacho, C., Coulouris, G., Avagyan, V., Ma, N., Papadopoulos, J., Bealer, K., & Madden, T. L. (2009). BLAST+: architecture and applications. BMC Bioinformatics, 10, 421. 10.1186/1471-2105-10-421

Celen, A. B., & Sahin, U. (2020). Sumoylation on its 25th anniversary: mechanisms, pathology, and emerging concepts. FEBS J, 287(15), 3110–3140. 10.1111/febs.15319

Clark, L., Sue-Ob, K., Mukkawar, V., Jones, A. R., & Sadanandom, A. (2022). Understanding SUMO-mediated adaptive responses in plants to improve crop productivity. Essays Biochem, 66(2), 155–168. 10.1042/EBC20210068

Contreras-Moreira, B., Saraf, S., Naamati, G., Casas, A. M., Amberkar, S. S., Flicek, P., Jones, A. R., & Dyer, S. (2023). GET_PANGENES: calling pangenes from plant genome alignments confirms presence-absence variation. Genome Biol, 24(1), 223. 10.1186/s13059-023-03071-z

Contreras-Moreira, B., Sharma, E., Saraf, S., Naamati, G., Gupta, P., Elser, J., Chebotarov, D., Chougule, K., Lu, Z., Wei, S., Olson, A., Tsang, I., Lodha, D., Zhou, Y., Yu, Z., Zhao, W., Zhang, J., Amberkar, S., Sue-Ob, K., … Jones, A. R. (2025). A pan-gene catalogue of Asian cultivated rice. bioRxiv.

Emenecker, R. J., Griffith, D., & Holehouse, A. S. (2021). Metapredict: a fast, accurate, and easy-to-use predictor of consensus disorder and structure. Biophys J, 120(20), 4312–4319. 10.1016/j.bpj.2021.08.039

Fernandes, T., Goncalves, N. M., Matiolli, C. C., Rodrigues, M. A. A., Barros, P. M., Oliveira, M. M., & Abreu, I. A. (2024). SUMOylation of rice DELLA SLR1 modulates transcriptional responses and improves yield under salt stress. Planta, 260(6), 136. 10.1007/s00425-024-04565-1

Fornasiero, A., Wing, R. A., & Ronald, P. (2022). Rice domestication. Curr Biol, 32(1), R20–R24. 10.1016/j.cub.2021.11.025

Garrido, E., Srivastava, A. K., & Sadanandom, A. (2018). Exploiting protein modification systems to boost crop productivity: SUMO proteases in focus. J Exp Bot, 69(19), 4625–4632. 10.1093/jxb/ery222

Ghimire, S., Tang, X., Zhang, N., Liu, W., Qi, X., Fu, X., & Si, H. (2020). Genomic Analysis of the SUMO-Conjugating Enzyme and Genes under Abiotic Stress in Potato (Solanum tuberosum L.). Int J Genomics, 2020, 9703638. 10.1155/2020/9703638

Gu, Z., Eils, R., & Schlesner, M. (2016). Complex heatmaps reveal patterns and correlations in multidimensional genomic data. Bioinformatics, 32(18), 2847–2849. 10.1093/bioinformatics/btw313

Hernández-Soto, A., Echeverría-Beirute, F., Abdelnour-Esquivel, A., Valdez-Melara, M., Boch, J., & Gatica-Arias, A. (2021). Rice breeding in the new era: Comparison of useful agronomic traits. Current Plant Biology, 27. 10.1016/j.cpb.2021.100211

Ibrahim, E. I., Attia, K. A., Ghazy, A. I., Itoh, K., Almajhdi, F. N., & Al-Doss, A. A. (2021). Molecular Characterization and Functional Localization of a Novel SUMOylation Gene in Oryza sativa. Biology (Basel), 11(1). 10.3390/biology11010053

Ingole, K. D., Dahale, S. K., & Bhattacharjee, S. (2021). Proteomic analysis of SUMO1-SUMOylome changes during defense elicitation in Arabidopsis. J Proteomics, 232, 104054. 10.1016/j.jprot.2020.104054

Jing, C. Y., Zhang, F. M., Wang, X. H., Wang, M. X., Zhou, L., Cai, Z., Han, J. D., Geng, M. F., Yu, W. H., Jiao, Z. H., Huang, L., Liu, R., Zheng, X. M., Meng, Q. L., Ren, N. N., Zhang, H. X., Du, Y. S., Wang, X., Qiang, C. G., … Ge, S. (2023). Multiple domestications of Asian rice. Nat Plants, 9(8), 1221–1235. 10.1038/s41477-023-01476-z

Jones, P., Binns, D., Chang, H. Y., Fraser, M., Li, W., McAnulla, C., McWilliam, H., Maslen, J., Mitchell, A., Nuka, G., Pesseat, S., Quinn, A. F., Sangrador-Vegas, A., Scheremetjew, M., Yong, S. Y., Lopez, R., & Hunter, S. (2014). InterProScan 5: genome-scale protein function classification. Bioinformatics, 30(9), 1236–1240. 10.1093/bioinformatics/btu031

Kim, D., Paggi, J. M., Park, C., Bennett, C., & Salzberg, S. L. (2019). Graph-based genome alignment and genotyping with HISAT2 and HISAT-genotype. Nat Biotechnol, 37(8), 907–915. 10.1038/s41587-019-0201-4

Kumar, S., Stecher, G., Li, M., Knyaz, C., & Tamura, K. (2018). MEGA X: Molecular Evolutionary Genetics Analysis across Computing Platforms. Mol Biol Evol, 35(6), 1547–1549. 10.1093/molbev/msy096

Kurepa, J., Walker, J. M., Smalle, J., Gosink, M. M., Davis, S. J., Durham, T. L., Sung, D. Y., & Vierstra, R. D. (2003). The small ubiquitin-like modifier (SUMO) protein modification system in Arabidopsis. Accumulation of SUMO1 and −2 conjugates is increased by stress. J Biol Chem, 278(9), 6862–6872. 10.1074/jbc.M209694200

Letunic, I., & Bork, P. (2024). Interactive Tree of Life (iTOL) v6: recent updates to the phylogenetic tree display and annotation tool. Nucleic Acids Res, 52(W1), W78–W82. 10.1093/nar/gkae268

Li, Y., Wang, G., Xu, Z., Li, J., Sun, M., Guo, J., & Ji, W. (2017). Organization and Regulation of Soybean SUMOylation System under Abiotic Stress Conditions. Front Plant Sci, 8, 1458. 10.3389/fpls.2017.01458

Lian, W., Zhan, X., Chen, D., Wu, W., Liu, Q., Zhang, Y., Cheng, S., Lou, X., Cao, L., & Hong, Y. (2024). Genome-Wide Identification and Characterization of the PPPDE Gene Family in Rice. Agronomy, 14(5). 10.3390/agronomy14051035

López-Fernández, H., Ferreira, P., Reboiro-Jato, M., Vieira, C. P., & Vieira, J. (2022). The pegi3s bioinformatics docker images project. Practical Applications of Computational Biology & Bioinformatics, 15th International Conference (PACBB 2021),

Mirdita, M., Schutze, K., Moriwaki, Y., Heo, L., Ovchinnikov, S., & Steinegger, M. (2022). ColabFold: making protein folding accessible to all. Nat Methods, 19(6), 679–682. 10.1038/s41592-022-01488-1

Orosa, B., Yates, G., Verma, V., Srivastava, A. K., Srivastava, M., Campanaro, A., De Vega, D., Fernandes, A., Zhang, C., Lee, J., Bennett, M. J., & Sadanandom, A. (2018). SUMO conjugation to the pattern recognition receptor FLS2 triggers intracellular signalling in plant innate immunity. Nat Commun, 9(1), 5185. 10.1038/s41467-018-07696-8

Paradis, E., & Schliep, K. (2019). ape 5.0: an environment for modern phylogenetics and evolutionary analyses in R. Bioinformatics, 35(3), 526–528. 10.1093/bioinformatics/bty633

Pei, W., Jain, A., Ai, H., Liu, X., Feng, B., Wang, X., Sun, Y., Xu, G., & Sun, S. (2019). OsSIZ2 regulates nitrogen homeostasis and some of the reproductive traits in rice. J Plant Physiol, 232, 51–60. 10.1016/j.jplph.2018.11.020

Pei, W., Jain, A., Sun, Y., Zhang, Z., Ai, H., Liu, X., Wang, H., Feng, B., Sun, R., Zhou, H., Xu, G., & Sun, S. (2017). OsSIZ2 exerts regulatory influences on the developmental responses and phosphate homeostasis in rice. Sci Rep, 7(1), 12280. 10.1038/s41598-017-10274-5

Pertea, M., Pertea, G. M., Antonescu, C. M., Chang, T.-C., Mendell, J. T., & Salzberg, S. L. (2015). StringTie enables improved reconstruction of a transcriptome from RNA-seq reads. Nature biotechnology, 33(3), 290–295.

Phosuwan, S., Nounjan, N., Theerakulpisut, P., Siangliw, M., & Charoensawan, V. (2024). Comparative quantitative trait loci analysis framework reveals relationships between salt stress responsive phenotypes and pathways. Front Plant Sci, 15, 1264909. 10.3389/fpls.2024.1264909

Rosa, M. T. G., Almeida, D. M., Pires, I. S., da Rosa Farias, D., Martins, A. G., da Maia, L. C., de Oliveira, A. C., Saibo, N. J. M., Oliveira, M. M., & Abreu, I. A. (2018). Insights into the transcriptional and post-transcriptional regulation of the rice SUMOylation machinery and into the role of two rice SUMO proteases. BMC Plant Biol, 18(1), 349. 10.1186/s12870-018-1547-3

Roy, D., & Sadanandom, A. (2021). SUMO mediated regulation of transcription factors as a mechanism for transducing environmental cues into cellular signaling in plants. Cell Mol Life Sci, 78(6), 2641–2664. 10.1007/s00018-020-03723-4

Sakai, H., Lee, S. S., Tanaka, T., Numa, H., Kim, J., Kawahara, Y., Wakimoto, H., Yang, C. C., Iwamoto, M., Abe, T., Yamada, Y., Muto, A., Inokuchi, H., Ikemura, T., Matsumoto, T., Sasaki, T., & Itoh, T. (2013). Rice Annotation Project Database (RAP-DB): an integrative and interactive database for rice genomics. Plant Cell Physiol, 54(2), e6. 10.1093/pcp/pcs183

Schrodinger, LLC. (2015). The AxPyMOL Molecular Graphics Plugin for Microsoft PowerPoint, Version 1.8.

Seok, H. Y., Bae, H., Kim, T., Mehdi, S. M. M., Nguyen, L. V., Lee, S. Y., & Moon, Y. H. (2021). Non-TZF Protein AtC3H59/ZFWD3 Is Involved in Seed Germination, Seedling Development, and Seed Development, Interacting with PPPDE Family Protein Desi1 in Arabidopsis. Int J Mol Sci, 22(9). 10.3390/ijms22094738

Srivastava, A. K., Zhang, C., Caine, R. S., Gray, J., & Sadanandom, A. (2017). Rice SUMO protease Overly Tolerant to Salt 1 targets the transcription factor, OsbZIP23 to promote drought tolerance in rice. Plant J, 92(6), 1031–1043. 10.1111/tpj.13739

Srivastava, A. K., Zhang, C., & Sadanandom, A. (2016). Rice OVERLY TOLERANT TO SALT 1 (OTS1) SUMO protease is a positive regulator of seed germination and root development. Plant Signal Behav, 11(5), e1173301. 10.1080/15592324.2016.1173301

Srivastava, A. K., Zhang, C., Yates, G., Bailey, M., Brown, A., & Sadanandom, A. (2016). SUMO Is a Critical Regulator of Salt Stress Responses in Rice. Plant Physiol, 170(4), 2378–2391. 10.1104/pp.15.01530

Sureshkumar, S., Bandaranayake, C., Lv, J., Dent, C. I., Bhagat, P. K., Mukherjee, S., Sarwade, R., Atri, C., York, H. M., Tamizhselvan, P., Shamaya, N., Folini, G., Bergey, B. G., Yadav, A. S., Kumar, S., Grummisch, O. S., Saini, P., Yadav, R. K., Arumugam, S., … Balasubramanian, S. (2024). SUMO protease FUG1, histone reader AL3 and chromodomain protein LHP1 are integral to repeat expansion-induced gene silencing in Arabidopsis thaliana. Nat Plants, 10(5), 749–759. 10.1038/s41477-024-01672-5

Teramura, H., Yamada, K., Ito, K., Kasahara, K., Kikuchi, T., Kioka, N., Fukuda, M., Kusano, H., Tanaka, K., & Shimada, H. (2021). Characterization of novel SUMO family genes in the rice genome. Genes Genet Syst, 96(1), 25–32. 10.1266/ggs.20-00034

Wang, H., Sun, R., Cao, Y., Pei, W., Sun, Y., Zhou, H., Wu, X., Zhang, F., Luo, L., Shen, Q., Xu, G., & Sun, S. (2015). OsSIZ1, a SUMO E3 Ligase Gene, is Involved in the Regulation of the Responses to Phosphate and Nitrogen in Rice. Plant Cell Physiol, 56(12), 2381–2395. 10.1093/pcp/pcv162

Yates, G., Srivastava, A. K., & Sadanandom, A. (2016). SUMO proteases: uncovering the roles of deSUMOylation in plants. J Exp Bot, 67(9), 2541–2548. 10.1093/jxb/erw092

Yuan, X., Luan, Y., Liu, D., Wang, J., Peng, J., Zhao, J., Li, L., Su, J., Xiao, Y., Li, Y., Ma, X., Zhu, X., Tan, L., Liu, F., Sun, H., Gu, P., Xu, R., Zhang, P., Zhu, Z., … Zhang, K. (2025). The SUMO-conjugating enzyme OsSCE1a from wild rice regulates the functional stay-green trait in rice. Plant Biotechnol J, 23(2), 615–631. 10.1111/pbi.14524

Zhang, S., Wang, S., Lv, J., Liu, Z., Wang, Y., Ma, N., & Meng, Q. (2018). SUMO E3 Ligase SlSIZ1 Facilitates Heat Tolerance in Tomato. Plant Cell Physiol, 59(1), 58–71. 10.1093/pcp/pcx160

Zhou, Y., Yu, Z., Chebotarov, D., Chougule, K., Lu, Z., Rivera, L. F., Kathiresan, N., Al-Bader, N., Mohammed, N., Alsantely, A., Mussurova, S., Santos, J., Thimma, M., Troukhan, M., Fornasiero, A., Green, C. D., Copetti, D., Kudrna, D., Llaca, V., … Wing, R. A. (2023). Pan-genome inversion index reveals evolutionary insights into the subpopulation structure of Asian rice. Nat Commun, 14(1), 1567. 10.1038/s41467-023-37004-y

